# A REM-active basal ganglia circuit that regulates anxiety

**DOI:** 10.1101/2023.07.27.550788

**Authors:** Wei Ba, Mathieu Nollet, Xiao Yu, Sara Wong, Andawei Miao, Esteban Beckwith, Edward C. Harding, Ying Ma, Raquel Yustos, Alexei L. Vyssotski, William Wisden, Nicholas P. Franks

**Affiliations:** Department of Life Sciences, Imperial College London, London SW7 2AZ, UK; UK Dementia Research Institute, Imperial College London, London SW7 2AZ, UK; Institute of Neuroinformatics, University of Zurich and ETH Zurich, Zurich 8057, Switzerland

**Keywords:** Basal ganglia, habenula, ventral tegmental area, rapid-eye-movement sleep, anxiety, defensive behavior

## Abstract

REM sleep has been hypothesized to promote emotional resilience, but any neuronal circuits mediating this have not been identified. We find that in mice, somatostatin (Som) neurons in the entopeduncular nucleus (EP^Som^)/internal globus pallidus are predominantly active selectively during REM sleep. This unique REM activity is necessary and sufficient for maintaining normal REM sleep. Inhibiting or exciting EP^Som^ neurons reduced or increased REM sleep duration, respectively. Activation of the sole downstream target of EP^Som^ neurons, Vglut2 cells in the lateral habenula (LHb), increased sleep via the ventral tegmental area (VTA). A simple chemogenetic scheme to periodically inhibit the LHb over 4 days selectively removed a significant amount of cumulative REM sleep. Chronic REM reduction correlated with mice becoming anxious and more sensitive to aversive stimuli. Therefore, we suggest that REM sleep, in part generated by the EP→LHb→VTA circuit identified here, could contribute to stabilizing reactions to habitual aversive stimuli.

## INTRODUCTION

The hypothesis that sleep serves a restorative function has focused largely on NREM sleep, the stage of sleep that occurs first [1]. Recently, however, REM sleep in humans has been found to be the deepest and most subjectively satisfying stage of sleep [2, 3], and is also the sleep state where capillary blood flow in the mouse brain is selectively boosted [4]. REM sleep has been hypothesized to promote emotional health or resilience [5-13], for example, in mice, by enabling contextual memory and/or forgetting about emotionally-salient stimuli [14-16], and diverse low-level chronic stressors substantially increases the amount of REM sleep [5, 17]. In these situations, more REM sleep could be an adaptation to deal with stress. Similarly, people living with severe depression or post-traumatic-stress disorder have elevated REM sleep [18], a potentially restorative mechanism [7]. An alternative interpretation is that more REM sleep in these conditions could be an abnormality contributing further to pathology [19]. Indeed, monoamine reuptake blockers, taken for depression, markedly decrease REM [20]. These apparent discrepancies as to whether changes in REM sleep are beneficial or harmful, remain unresolved.

Although the brainstem circuitry generating atonia during REM is well understood [21], knowledge of how REM is generated in the forebrain circuitry is incomplete [22-24]. Previously we found that genetically blocking glutamate release from mouse lateral habenula cells produces severe sleep-wake fragmentation [25], potentially revealing a new sleep-wake pathway. Here, starting by looking at upstream and downstream partners of LHb cells, we found cells in the basal ganglia (the EP nucleus, also known as the habenular-projecting globus pallidus or GPh, or the internal globus pallidum in primates) projecting to the LHb and then to the VTA, that induce NREM and REM sleep, mapping onto previous circuitry that gets activated with aversion, disappointment, learned helplessness and anxiety [26-29]. Because of this overlap, we suggest that REM sleep contributes to emotional stability, and test this hypothesis by selectively reducing REM sleep originating from this circuit.

## RESULTS

### EP^Som^ neurons are strongly REM sleep active

The LHb receives diverse inputs [30-35], but including a unique projection from somatostatin (Som)-glutamate-GABA neurons in the entopeduncular nucleus [26, 36-38]. We performed anterograde tracing by injecting *AAV-DIO-mCherry* in the EP of *Som-Cre* mice and confirmed that EP^Som^ neurons project exclusively to the LHb as previously reported [26, 36, 39-41] (Figure 1A and 1B). We then used calcium photometry to study the natural dynamics of the EP^Som^ neurons across the sleep-wake cycle. *AAV-DIO-GCaMP6s* was injected into the EP of *Som-Cre* mice with an optical fiber implanted over the EP to acquire population activity of EP^Som^ neurons (Figure 1C and Figure S1A). We observed the highest activity of EP^Som^ neurons during REM sleep, while the activity during NREM sleep and wakefulness was suppressed (Figure 1D-F). The baseline Ca^2+^ transients began to elevate at the end of NREM episodes and we observed a time-locked increase (*P* = 0.026; *n* = 6) at the transition from NREM to REM. Conversely, the fluorescence signal decreased (*P* = 0.022; *n* = 6) during REM-WAKE transitions. Recordings from EP^Som^-GFP control mice displayed no such vigilance state-dependent variations (Figure S1B and S1C).

**Figure 1.**
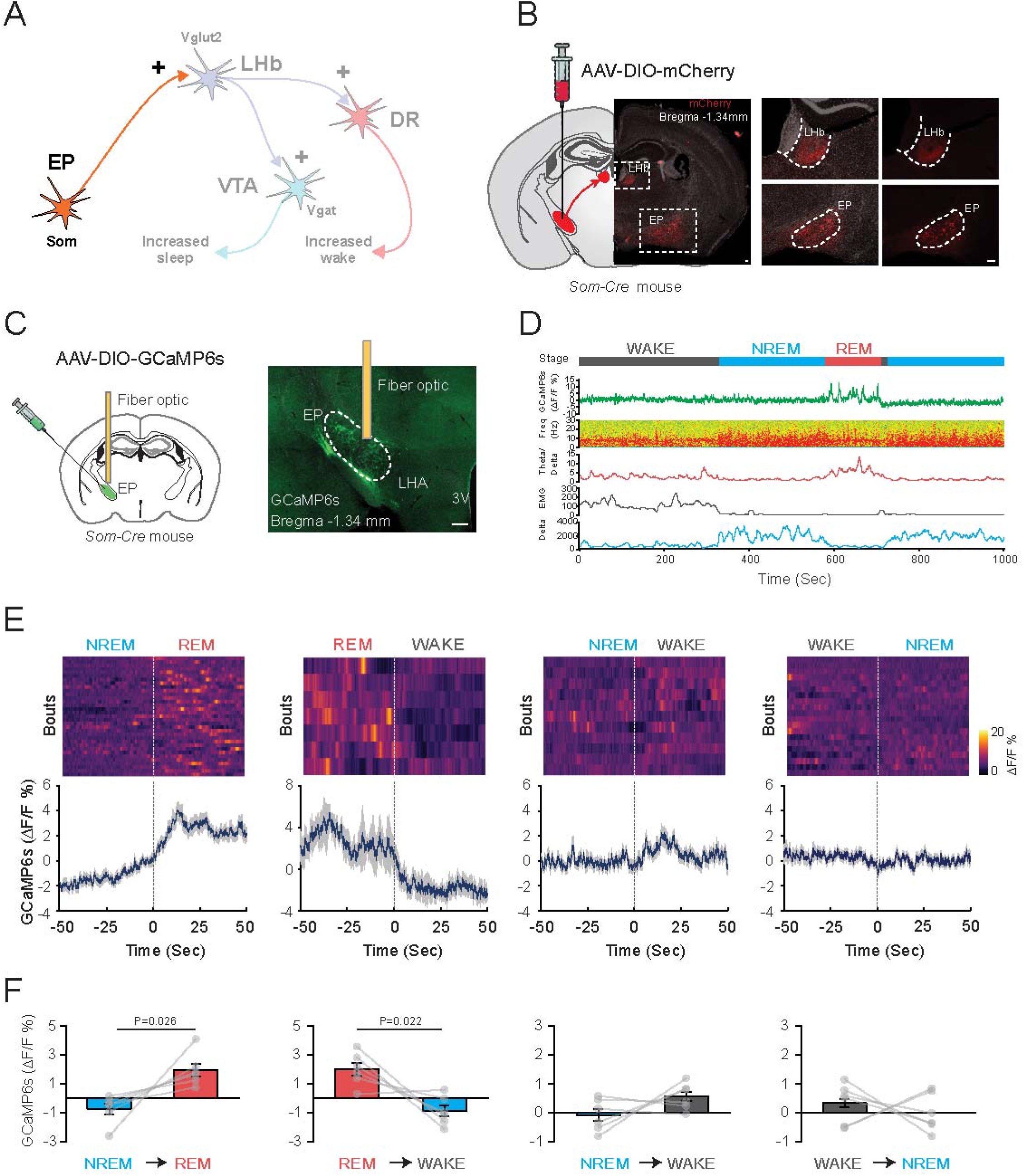
Spontaneous activity of EP^Som^ neurons across sleep-wake states. (A) Conceptual schematic diagram of the neuronal circuit examined in this figure. EP: Entopeduncular nucleus, LHb: Lateral habenula, DR: Dorsal raphe nucleus, VTA: Ventral tegmental area. (B) Anterograde tracing confirms that EP^Som^ neurons project exclusively to the LHb. Representative images of the cell bodies in the EP and terminals in the LHb after *AAV-DIO-mCherry* was injected into the EP of *Som-Cre* mice. Left, a coronal section includes both EP and LHb (indicated in dashed boxes). Right, mCherry expressing neurons in the EP and its projections in the LHb with higher magnification. No projections in other brain areas were observed. Grey, nuclei stained with Hoechst 33342; Red, mCherry, Scale bars, 100 μm. (C) Fiber photometry with EEG/EMG recordings in freely moving mice. *AAV-DIO-GCaMP6s* was injected into the EP of *Som-Cre* mice and a fiber optic was implanted above the EP to record activity (scale bar, 200 μm). LHA, lateral hypothalamic area, 3V, third ventricle. (D) Representative fiber photometry recording aligned with the EEG spectra and EMG recordings during wakefulness, NREM and REM sleep. (E) Combined fluorescence responses during transitions between vigilance states for all mice (*n* = 6). Top, individual traces with color-coded fluorescence intensity. Bottom, mean responses of each transition (blue) ± SEM (grey). (F) Summary of transitions between states. P values are given when <0.05.

### EP^Som^ neurons bidirectionally regulate REM sleep

To assess whether EP^Som^ activity is necessary for natural REM sleep, we inhibited these neurons chemogenetically. *AAV-DIO-hM4Di-mCherry* (referred to as “hM4Di” hereafter) was bilaterally injected into the EP of *Som-Cre* mice (Figure 2A and S2A). Our preliminary tests indicated a sleep-suppressing trend when inhibiting EP^Som^ neurons, we further evaluated this effect during the lights-on phase when mice are mostly inactive and show abundant sleep. Systemic administration of 1 mg/kg clozapine-*N*-oxide (CNO) at the beginning of the lights-on phase significantly (*P* = 0.0004; *n* = 7) reduced the total duration of REM sleep during the 3 hours following CNO injection, as compared with saline-injected controls (Figure 2B and 2C). Reduced REM sleep amounts were due to decreased numbers of long (>1-min) REM-sleep episodes (from 54% to 29% *P* = 0.03; n = 7). In contrast, total duration of wakefulness and NREM sleep remained unchanged (Figure 2B and 2C). Furthermore, chemogenetic inhibition of EP^Som^ neurons significantly (*P* = 0.02; *n* = 7) increased the latency to REM sleep without affecting NREM latency (Figure 2C). We next selectively lesioned EP^Som^ cells using *AAV-DIO-CASP3* injected into the EP of *Som-Cre* mice. Four weeks after the lesions, shortened times in REM sleep were also observed (Figure 2D). Following ablation (79% ablation; *P* = 0.009, Figure S2B), the total duration of REM sleep in the light phase consistently (*P* = 0.0004; *n* = 8 control and *n* = 11 caspase) declined, further confirming that these neurons contribute to normal REM-sleep structure. This effect was selectively confined to the 12-hour light part of the daily cycle.

**Figure 2.**
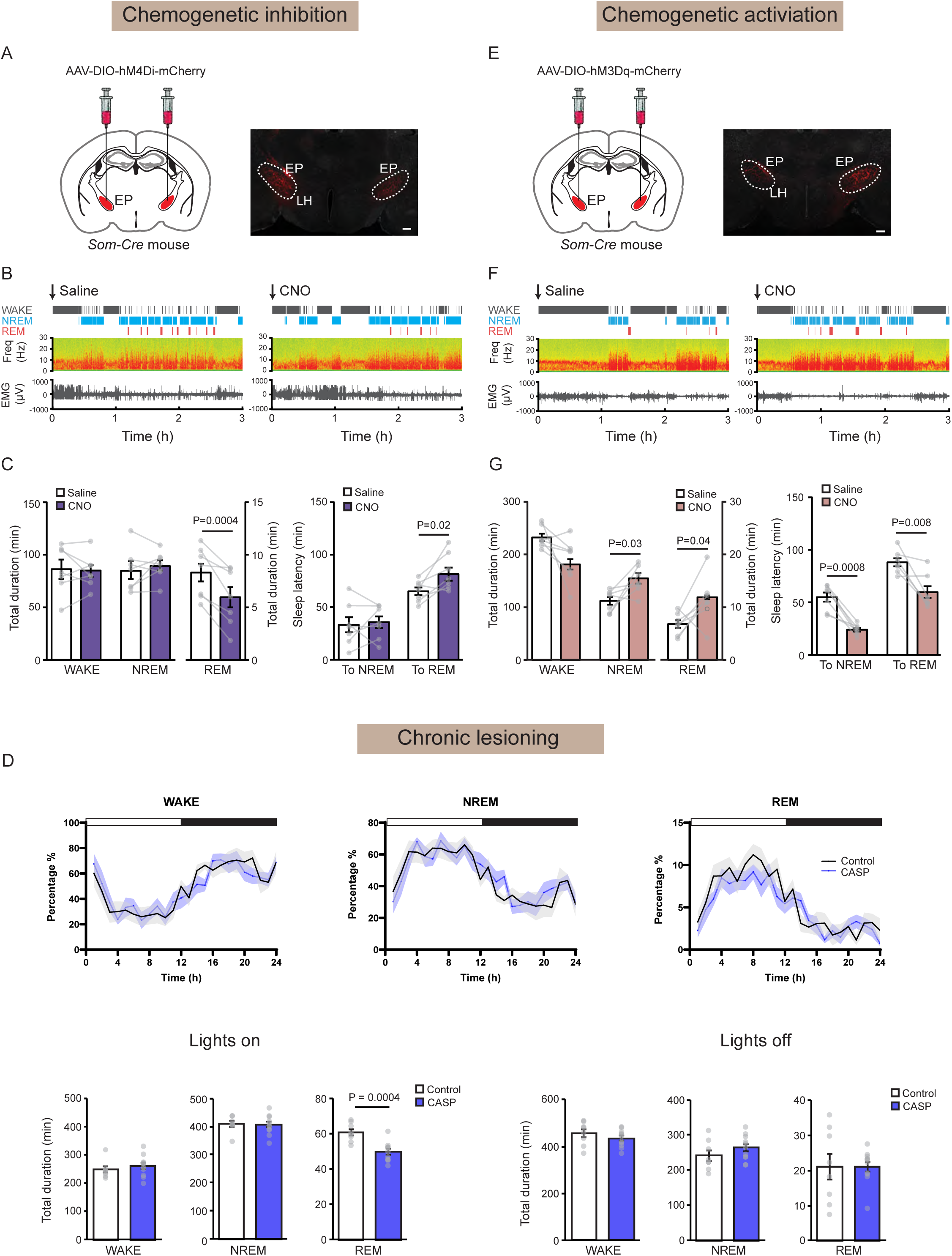
EP^Som^ neurons are sufficient and necessary for normal REM sleep. (A-C) Chemogenetic inhibition of EP^Som^ neurons suppresses REM sleep. (A) Schematic illustrating inhibition of EP^Som^ neurons using DREADD receptors. *AAV-DIO-hM4Di-mCherry* was injected into the EP of *Som-Cre* mice. Immunostaining confirmed the expression of hM4Di-mCherry in the EP. LH, lateral hypothalamus; Scale bar, 200 μm. (B) Representative EEG/EMG recordings post-saline or CNO (1 mg/kg) injections. (C) Summarized changes on sleep duration and latency. Data are means ± SEMs (*n* = 7). (D) Sleep analysis of control and CASP3 mice. Top, 24h sleep recording post-saline (black) or CNO (blue) injection. Bottom, quantification of each vigilance state in the lights on or lights off phases. Chronic lesioning of EP^Som^ neurons reduced REM sleep duration in the lights on phase without affecting NREM and wakefulness (*P* = 0.0004, control *n* = 8, CASP *n* = 11). (E-G) Chemogenetic activation of EP^Som^ neurons increases sleep. (E) Schematic of activation of EP^Som^ neurons using DREADD receptors. *AAV-DIO-hM3Dq-mCherry* was injected into the EP. (F) Representative traces of EEG/EMG recordings post-saline or CNO injections. (G) Summarized changes of sleep duration and latency. Data are means ± SEMs (*n* = 7).

To examine whether EP^Som^ neurons can elicit REM sleep in fully awake mice, we expressed *AAV-DIO-hM3Dq-mCherry* (referred to as “hM3Dq” hereafter) in the EP^Som^ neurons (Figure 2E and S3A) and performed intraperitoneal injections of saline or CNO in the dark period when mice show high levels of arousal. Compared with the saline control group, 1 mg/kg CNO injection increased the (*P* = 0.0049; *n* = 2 control and *n* = 3 hM3Dq) number of c-FOS positive neurons in the EP (Figure S3B) and induced NREM and REM sleep (Figure 2F and 2G). The total duration of both NREM and REM sleep significantly (*P* = 0.03 and *P* = 0.04 respectively; *n* = 7) increased during the 6 hours post CNO injection (Figure 2F and 2G). The latencies to both NREM and REM sleep were reduced (*P* = 0.0008 and *P* = 0.008 respectively; *n* = 7) by CNO (Figure 2G). Given that EP^Som^ neurons are silent during NREM sleep, and that REM sleep only occurs following NREM sleep in physiological conditions, we interpret the increased NREM sleep as a necessary element for inducing REM sleep.

### EP^Som^ neurons regulate REM sleep via their sole target - the LHb

We reasoned that EP^Som^ neurons regulate REM sleep via their sole downstream target, the LHb. Three experiments were carried out to confirm this. First, we examined whether the output from EP to LHb correlated with REM sleep by monitoring the Ca^2+^ transients at EP→LHb terminals. We injected *AAV-DIO-GCaMP6s* in EP^Som^ neurons and positioned the optical fiber over the LHb. Our results showed that as for the EP^Som^ somata, the Ca^2+^ transients at EP→LHb terminals peaked during REM sleep (Figure S4). NREM-REM sleep transitions were associated with prominent increases of activity at these terminals (*P* = 0.0002). Next, we examined whether the LHb is similarly modulated during REM sleep by measuring the Ca^2+^ transients in the cell bodies of LHb neurons across the sleep-wake cycle (Figure 3A and 3B). (Note: we used *Vglut2* gene expression as a local selective marker for lateral habenula cells, so as not to get contaminating expression from the overlying hippocampal dentate granule cells). We found that LHb^Vglut2^ neurons were most active during wakefulness and REM sleep and suppressed during NREM sleep (Figure 3C-E). The Ca^2+^-induced fluorescence increased significantly at NREM to REM transitions. Conversely, LHb^Vglut2^ activity decreased when animals entered NREM sleep. Interestingly, the activity in the LHb during wakefulness is attributed, at least in part, to its excitatory input from the lateral preoptic area (LPO) (Figure S5). Finally, we examined whether the LHb^Vglut2^ neurons can bidirectionally regulate REM sleep. We expressed hM4Di in LHb^Vglut2^ neurons and performed EEG/EMG analysis comparing control saline injections with CNO injections (Figure 4A). CNO-mediated inhibition of LHb^Vglut2^ neurons decreased REM sleep, leaving NREM sleep and wakefulness intact (Figure 4B and 4C). This reduction was due to fewer REM episodes (Figure S6A). Conversely, chemogenetically activating LHb^Vglut2^ neurons prolonged both NREM and REM sleep at the expense of wakefulness (Figure 4D-F). Increased episode number and duration of REM sleep were observed (Figure S6B). In control experiments, CNO (1 mg/kg; *i.p.*) had no effect on vigilance states in *Vglut2-Cre* mice (Figure S7). Taken together, these findings reveal a close correlation of LHb activity with REM sleep as well as a capacity of the LHb for the bidirectional regulation of REM sleep, thereby supporting the idea that the EP→LHb circuit contributes to maintaining normal REM sleep.

**Figure 3.**
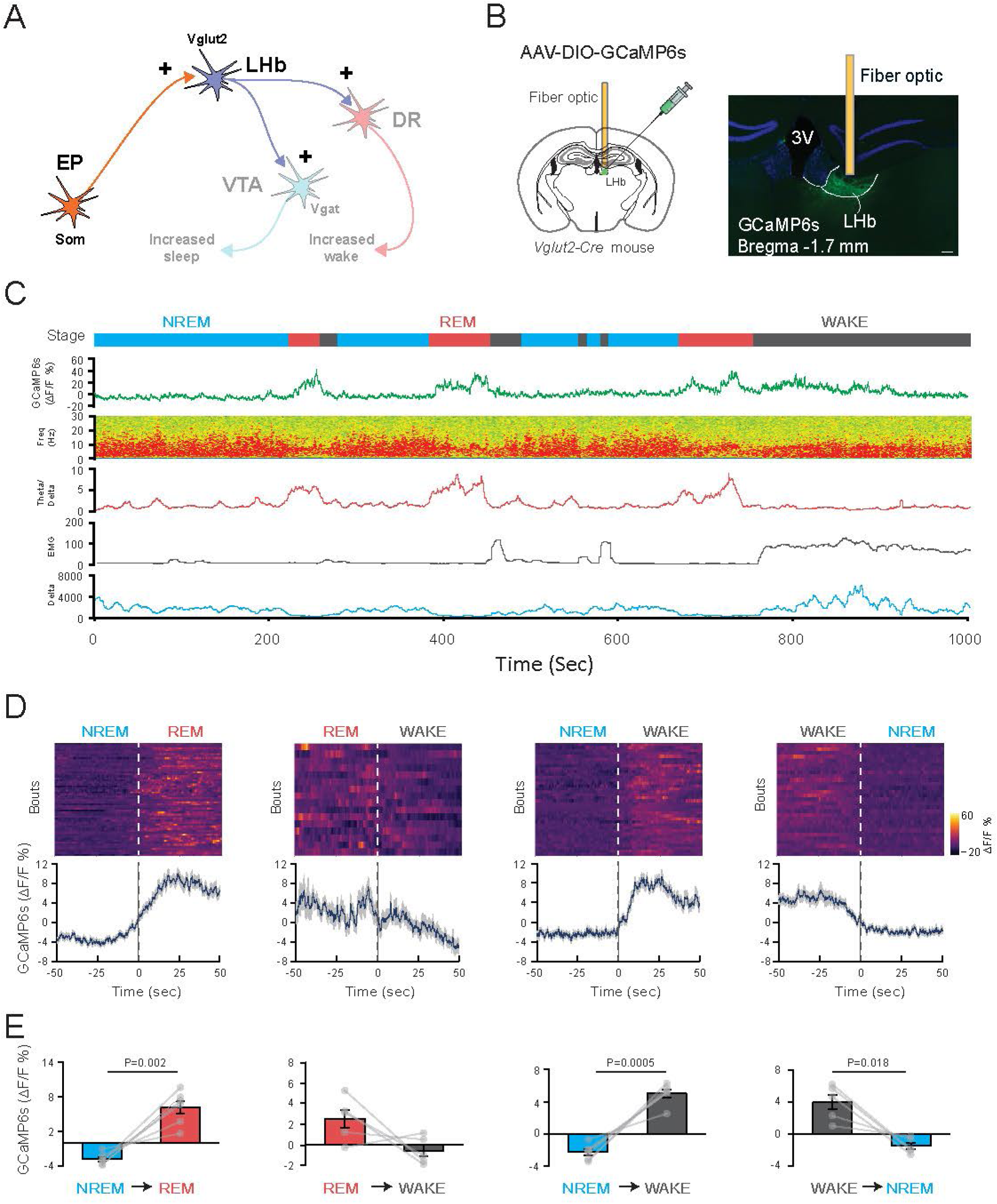
Spontaneous activities of LHb^Vglut2^ neurons across sleep-wake. (A) Conceptual schematic circuit diagram. EP: Entopeduncular nucleus, LHb: Lateral habenula, DR: Dorsal Raphe nucleus, VTA: Ventral tegmental area. (B) Fiber photometry with EEG/EMG recordings in freely moving mice. Left, schematic illustration of experimental setup; Right, *AAV-DIO-GCaMP6s* was injected into the LHb of *Vglut2-Cre* mice and a fiber optic was implanted above the LHb to record activity; 3V, third ventricle (scale bar, 100 μm). (C) Representative Ca^2+^ signals aligned with the EEG spectra and EMG recordings during wakefulness, NREM and REM sleep. (D) Combined fluorescence responses during transitions between vigilance states for all mice (*n* = 6). Top, individual traces with color-coded fluorescence intensity. Bottom, mean responses of each transition (blue) ± SEM (grey). (E) Summary of transitions between states. P values are given when <0.05.

**Figure 4.**
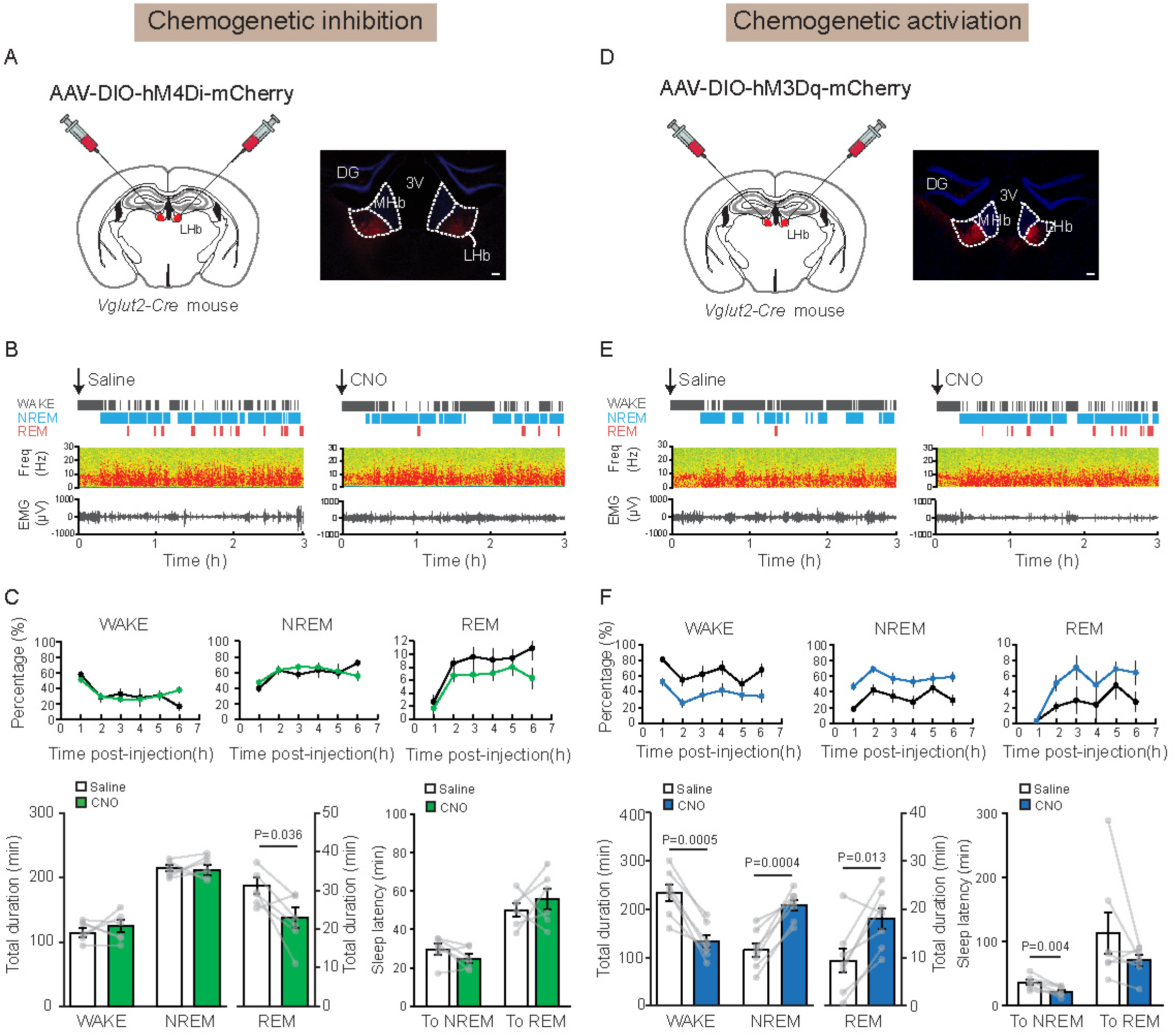
Casual effects of LHb^Vglut2^ neurons on REM sleep. (A to C) Chemogenetic inhibition of LHb^Vglut2^ neurons suppresses REM sleep. (A) Schematic illustrating inhibition of LHb^Vglut2^ neurons using DREADD receptors. *AAV-DIO-hM4Di-mCherry* was injected into the LHb of *Vglut2-Cre* mice. Immunostaining confirmed the expression of hM4Di-mCherry in the LHb; DG, dentate granule cells; MHb, medial habenula; 3V, third ventricle; (B) Representative EEG/EMG recordings post-saline or CNO (1 mg/kg) injection. (C) Summarized changes on sleep duration and latency. Data are means ± SEMs (*n* = 7). (D to F) Effects of chemogenetic activation of LHb^Vglut2^ neurons on sleep. (D) Schematic illustrating activation of LHb^Vglut2^ neurons using DREADD receptors. *AAV-DIO-hM3Dq-mCherry* was injected into the LHb of *Vglut2-Cre* mice. (E) Representative traces of EEG/EMG recordings in the saline or CNO-injected groups. (F) Summarized changes of sleep duration and latency. Data are means ± SEMS (*n* = 7).

### LHb→VTA terminals are most active during REM sleep and its selective activation promotes REM sleep

We next sought to identify the output pathway from the LHb mediating the REM-sleep-regulating effect. Previous studies identified multiple synaptic targets of the glutamatergic LHb neurons. We focused on the dorsal raphe nucleus (DRN) and the ventral tegmental area (VTA) [30], two areas modulating motivation/aversion and previously identified as contributing to sleep-wake regulation [42, 43]. *In-vivo* fiber-photometry recordings at LHb→DRN and LHb→VTA terminals across sleep-wake cycle were achieved similarly as described above, by expressing *AAV-DIO-GCaMP6s* in LHb^Vglut2^ neurons, and with optical fibers placed over terminals in either the DRN or VTA (Figure 5A). We observed a wake-specific calcium signal at the LHb→DRN terminals, while the activity was absent during sleep (Figure 5B and S8); indeed, the LHb→DRN projection has been found to induce aggressive arousal [44]. In contrast, NREM-REM transitions were consistently associated with prominent increases in the activity of LHb→VTA terminals (Figure 5C-E). To further establish whether the LHb promotes sleep through its projections to the VTA, we expressed hM3Dq in the LHb, and unilaterally delivered CNO, via a cannula, into the VTA to excite the LHb glutamatergic terminals (Figure S9A). Infusing CNO (0.2 µl of 3 µM, over 5 minutes) into the VTA elicited both NREM and REM sleep, recapitulating the effects of activating the LHb somata (Figure S9B-C).

**Figure 5.**
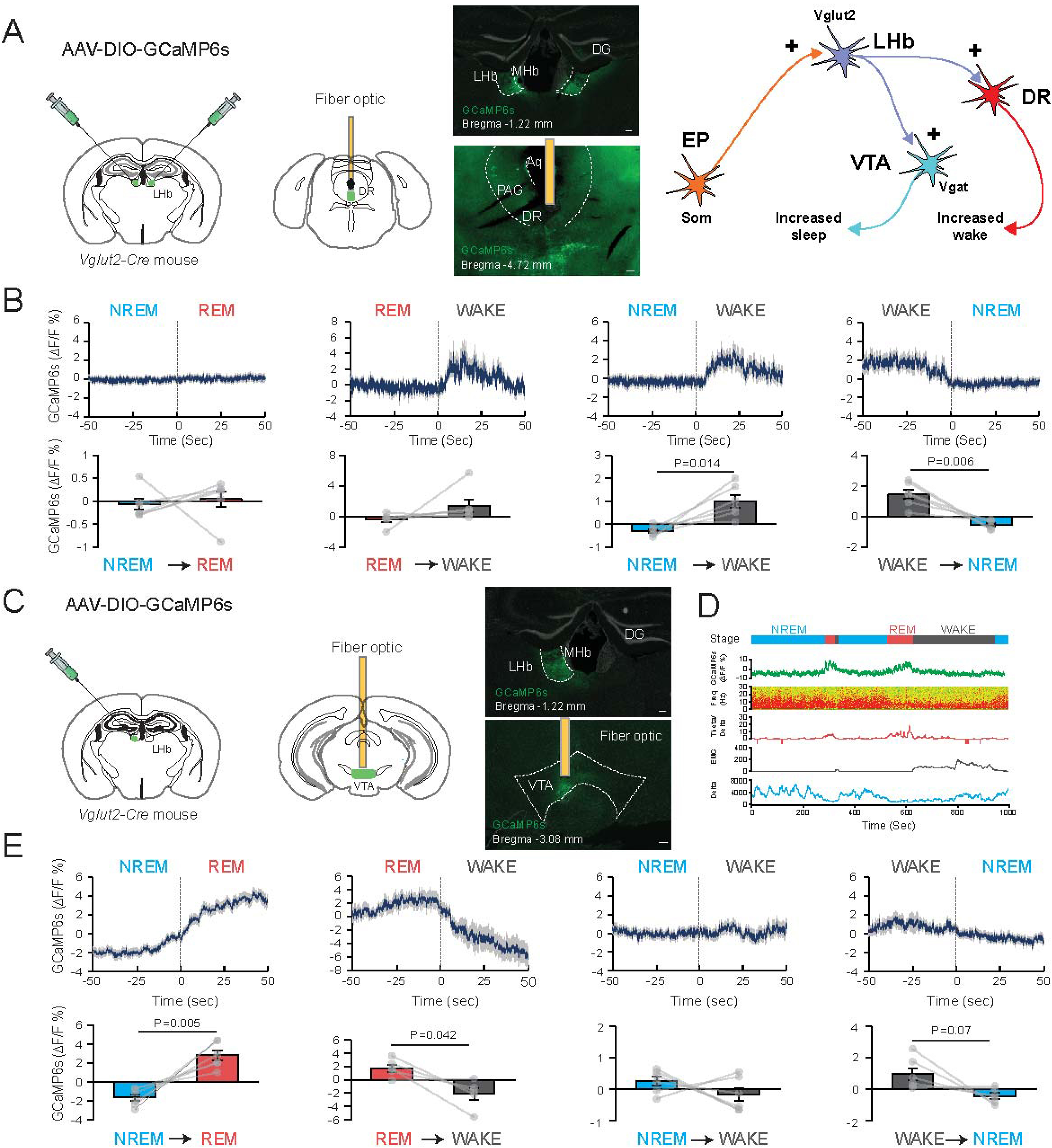
Two downstream targets of the LHb separately drive WAKE and REM. (A) Spontaneous activity across sleep-wake states measured by fiber photometry of LHb terminals in the dorsal raphe (DR) nucleus. Left, schematic illustration of experimental setup; *AAV-DIO-GCaMP6s* was injected into the LHb of *Vglut2-Cre* mice and a fiber optic was implanted above the DR to record terminal activity. Aq, ventricle; DG, dentate granule cells; MHb, medial habenula; PAG, periaqueductal grey (scale bar, 100 μm). Right, conceptual schematic circuit diagram. EP: Entopeduncular nucleus, LHb: Lateral habenula, DR: Dorsal Raphe nucleus, VTA: Ventral tegmental area. (B) Combined fluorescence responses during transitions between vigilance states for all mice (*n* = 6). Mean responses of each transition (blue) ± SEM (grey). *P* values are given when <0.05. (C) Spontaneous activity measured by fiber photometry across sleep-wake states of LHb terminals in the VTA. Left, schematic illustration of experimental setup; center, *AAV-DIO-GCaMP6s* was injected into the LHb of *Vglut2-Cre* mice and a fiber optic was implanted above the VTA to record activity; DG, dentate granule cells; MHb, medial habenula (scale bar, 100 μm). Right, conceptual schematic circuit diagram. EP: Entopeduncular nucleus, LHb: Lateral habenula, DR: Dorsal Raphe nucleus, VTA: Ventral tegmental area. (D) Representative traces of Ca^2+^ signals aligned with EEG and EMG recordings during WAKE, NREM and REM sleep indicated with grey, cyan and red respectively. (E) Combined fluorescence responses during transitions between vigilance states for all mice (*n* = 6). Mean responses of each transition (blue) ± SEM (grey). *P* values are given when <0.05. (F to H) Local chemogenetic activation of LHb^Vglut2^ terminals in the VTA neurons increases REM and NREM sleep. (F) Excitation of LHb^Vglut2^ terminals in VTA neurons using hM3Dq-mCherry receptors. *AAV-DIO-hM3Dq-mCherry* was injected into the LHb of *Vglut2-Cre* mice. Immunostaining confirmed the expression of hM3Dq-mCherry in the LHb soma and terminals in the VTA. (Scale bars, 100 μm.) (G) Representative EEG/EMG recordings post-saline or CNO infusion into the VTA. (H) Summarized changes on WAKE, NREM and REM duration. Data are means ± SEM (*n* = 6).

### Chronic REM sleep restriction increases anxiety level and alters defensive behavior

We assessed the functional significance of REM sleep generated by the circuit. We discovered that repetitive dosing of LHb*^Vglut2-hM4Di^* mice with CNO led to a chronic reduction of REM sleep. LHb*^Vglut2-hM4Di^* mice were given saline or CNO once a day at the beginning of the inactive lights-ON phase for consecutive four days. This reduced REM sleep by 24%, 15%, 11% and 7% over the four days but did not change the measurable amount of NREM sleep or WAKE (Figure S10). Then, 16 hours after the last saline or CNO administration on day 4, mice were evaluated in several behavioral tests (Figure 6A). In the light-dark test (Figure 6B), mice with reduced REM sleep increased their time in the dark compartment with a corresponding decrease in the light chamber. General locomotion of the mice injected with saline or CNO remained unchanged, as indicated by the same amount of entry numbers into the chambers (Figure 6B). Similarly, compared with control mice, mice spent less time in the open arm in an elevated plus maze test when REM sleep was suppressed (Figure 6C). These data showed that shortened REM sleep induced anxiety-like behaviors in mice.

**Figure 6.**
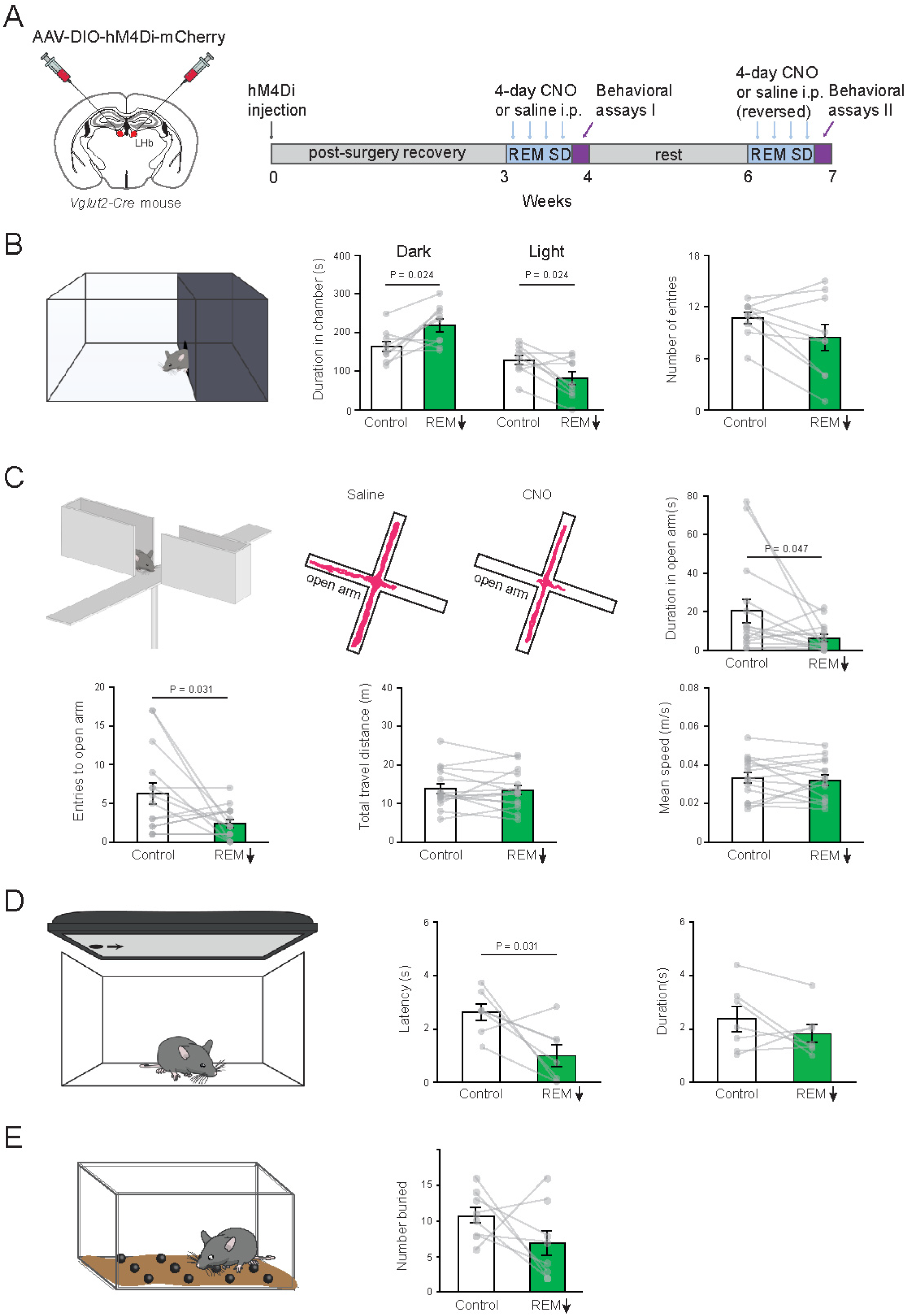
Chronic and selective reduction in REM sleep is accompanied by increased anxiety and altered defensive behavior. (A) Schematic illustration of experimental setup. Three weeks following transduction of LHb neurons in *Vglut2-Cre* mice with *AAV-DIO-hM4Di-mCherry*, one systemic injection of CNO (1mg/ml *i.p.*), or saline, each day for 4 days caused a cumulative reduction in REM (see Figure S10). Behavioral assays were carried out 16 hours after the last saline or CNO administration on day 4 following the REM-reduction protocol. After two weeks of rest, saline, or CNO (1mg/ml *i.p.*), was then again injected *i.p.* each day for 4 days followed by a repeat of the behavioral test. (B) Light/Dark box assay. Following REM reduction, mice chose to spend significantly more time in the dark region, compared to the light region. The motor activity, as assessed by the number of entries was unchanged. (C) In an elevated maze, following REM reduction, mice spent significantly less time in the open arm of the maze, with no change in motor activity. Typical trajectories for a control mouse and one with chronic REM reduction. Mice with chronic REM reduction spent less time and had fewer entries into the open part of the maze, but the total distance traveled, and the average speed did not change. (D) When presented with a potential looming threat (a black disk moving above them), mice responded by freezing (after a latency interval) and remaining immobile for a certain duration. Following the REM reduction protocol, the time before freezing (latency) reduced significant reduction in latency, but with no significant change in the duration of immobility. (E) REM sleep diminishment did not influence how many marbles mice buried.

To further assess whether REM sleep contributes to the performance of innate behaviors, such as defensive behaviors, we performed a looming test in which a moving disk is presented to mimic distal threats. In this test, mice generally respond with freezing behaviors [45]. When REM sleep was restricted for four days, CNO-treated LHb*^Vglut2-hM4Di^* mice responded to the stimuli much faster than the control group, indicated by reduced latencies to freeze (Figure 6D). The duration of freezing behaviors was unaltered, however (Figure 6D). Thus, REM sleep deficits induced sensitization of defensive responses to visual threats. We observed no change in the marble burying test (*P* = 0.2; *n* = 8) (Figure 6E), indicating that reducing REM sleep did not induce repetitive behaviors in mice. Overall, the behavioral results show that baseline REM sleep could contribute to stabilizing behavioral reactions to habitual aversive stimuli.

## DISCUSSION

The pathways that contribute to REM sleep generation are still being elucidated [21-24, 46-49]. The basal ganglia, a collection of subcortical nuclei that include the nucleus accumbens, the caudate-putamen, the globus pallidus externa, the EP (globus pallidus interna), and the substantia nigra, have all been implicated in the induction of NREM sleep or wakefulness, depending on the specific cell type [50-53]. By contrast, the strongly REM-active nature of the EP (Som-expressing) cells in basic home cage conditions of the mice are a striking feature of our findings. These EP^Som^ cells are unusual in that in both primates and rodents, they project to just one target, the LHb [37], a nucleus that we previously showed is needed for consolidated sleep [25]. The same circuitry of EP^Som^ cells and their downstream target, the LHb cells, fires with aversive events or disappointment [26, 28, 29], in other words, when an outcome is worse than expected [30, 54]. This EP→LHb→VTA circuit helps animals learn about negative experiences and/or adopt passive coping strategies [54-56]. That this circuitry seems to have an exact mapping with the REM sleep circuitry we report here seems more than a coincidence. Empirically, we discovered that a simple chemogenetic scheme to periodically chemogentically inhibit the LHb over 4 days selectively removed a significant amount of cumulative REM sleep, without affecting measurable amounts of NREM or wakefulness. Testing 16 hours later after the last inhibitory dose, chronic REM reduction correlated with mice becoming more sensitive to aversive stimuli. Therefore, we suggest that baseline REM sleep could contribute to stabilizing reactions to habitual aversive stimuli.

The LHb is a relay nucleus with many channels (reflecting diverse inputs), and receives innervations from about forty brain regions, including from the EP, but also from the hypothalamus, midbrain and brainstem areas [30-33, 35]. Our data showed that the input from the preoptic area of the hypothalamus is selectively active during wakefulness, likely contributing to the wake-active calcium signals in the LHb we observed. All subtypes of LHb cell release glutamate onto distal targets, and a few likely co-release GABA and glutamate [33]. We found that the outputs to the dorsal raphe are not sleep-active; but the LHb terminals in the VTA are, and could induce sleep by activating GABA neurons, so potentially feeding into a circuit we identified for how certain stressors (social defeat stress) induce sleep [57]. In social defeat stress, hypothalamic and brainstem stress pathways activate VTA GABA cells to promote sleep [57]. But milder stress and habitual aversive stimuli could activate the VTA sleep circuitry via the EP^Som^→LHb branch to regulate sleep. A further feature is that EP^Som^ cells co-release both GABA and glutamate, and so can simultaneously excite via ionotropic glutamate receptors, and inhibit via GABA_A_ receptors the postsynaptic LHb cells [38]. Based on our results, the glutamate component would be predicted to enhance REM and the GABA component decrease REM sleep. Certainly, our results indicate that the EP^Som^ cells predominately excite the LHb, consistent with previous reports [36]. But given that the balance of glutamate and GABA transmission at EP^Som^ to LHb synapses could vary with different behaviors, motivations and stressors [38, 58, 59], certain stressors and experiences could enhance REM sleep, whereas others could decrease it.

An overlap exists between the consequences of activating the EP→LHb→VTA sleep circuit we have identified and the state of depression. Non-medicated depressed patients often enter REM sleep more quickly, and their REM lasts longer [18]. Moreover, the LHb is more active during depression [60], and we predict that it is this habenula overactivity that causes the enhanced REM in this condition. The enhanced REM in people living with depression is often assumed to be an adverse outcome. However, this enhanced REM could, in fact, be beneficial and aid emotional processing, rather than being harmful. Indeed, this logic suggests that drugs or other treatments that selectively enhance REM might be beneficial in treating depression (current drugs suppress REM sleep [20]). Thus, understanding the sleep-promoting aspects of the EP→LHb→VTA sleep-promoting circuitry could allow insights to improve mental health.

## STAR★Methods

### Key resources table

STAR★Methods
| REAGENT or RESOURCE | SOURCE | IDENTIFIER |
| --- | --- | --- |
| **Antibodies** |  |  |
| c-Fos | Sigma-Aldrich | ABE457 |
| rat monoclonal mCherry | Thermo Fisher Scientific | Cat# M11217 |
| chicken polyclonal GFP | Abcam | Cat# ab13970 |
| Alexa Fluor 488 goat anti-rabbit | Invitrogen | Cat# A11108 |
| Alexa Fluor 488 goat anti-mouse | Invitrogen | Cat# A11001 |
| Alexa Fluor 594 goat anti-rabbit | Invitrogen | Cat# A11037 |
| Alexa Fluor 594 goat anti-mouse | Invitrogen | Cat# A11005 |
| Alexa Fluor 594 goat anti-rat | Invitrogen | Cat# A11007 |
| **Bacterial and virus strains** |  |  |
| Adenovirus helper plasmid *pFΔ6* | Donated by M Klugmann | NA |
| AAV helper plasmid *pH21* (AAV1) | Donated by M Klugmann | NA |
| AAV helper plasmid *pRVI*(AAV2) | Donated by M Klugmann | NA |
| *pAAV-hSyn-DIO-hM3Dq-mCherry* | Gift from Bryan L. Roth | Addgene plasmid 44361 |
| *pAAV-hSyn-DIO-hM4Di-mCherry* | Gift from Bryan L. Roth | Addgene plasmid 44362 |
| *pAAV-hSyn-DIO-GCaMP6s* | Addgene | Addgene plasmid 184284 |
| *pGP-CMV-GCaMP6s* | Addgene | Addgene plasmid 40753 |
| *pAAV-EF1α-DIO-taCASP3-TEV* | Gift from Nirao Shah | Addgene plasmid 45580 |
| **Chemicals, peptides, and recombinant proteins** |  |  |
| clozapine-N-oxide | Sigma-Aldrich | C0832 |
| Hoechst 33342 | Thermo Fisher Scientific | H3507 |
| **Experimental models: Organisms/strains** |  |  |
| Mouse: *Vglut2-ires-Cre* (*(Slc17a6<sup>tm2(cre)Lowl</sup>/J)*) | Jackson Lab | JAX Stock 016963 |
| Mouse: *Vgat-ires-Cre* (*(Slc32a1<sup>tm2(cre)Lowl</sup>/J)*) | Jackson Lab | JAX Stock 016962 |
| Mouse: *Som-ires-Cre* (*(Sst<sup>tm2.1(cre)Zjh</sup>/J)*) | Jackson Lab | JAX Stock 013044 |
| **Software and algorithms** |  |  |
| Spike 2 | Cambridge Electronic Design Limited | https://ced.co.uk/products/spkovin |
| Matlab | MathWorks | https://uk.mathworks.com/ |
| Fiji | Fiji | https://fiji.sc/ |

### Resource Availability

#### Lead contact

Further information and requests for resources and reagents should be directed to and will be fulfilled by the lead contact and corresponding authors: Nicholas P. Franks or William Wisden.

#### Materials availability

This study did not generate new unique reagents.

### Experimental model and subject details

All procedures were approved by the Animal Welfare Ethical Review Body at Imperial College London. Experiments were performed in accordance with the UK Home Office Animal Procedures Act (1986). *Vglut2-ires-Cre* mice (*Slc17a6^tm2(cre)Lowl^/J*) and *Vgat-ires-Cre* mice (*Slc32a1^tm2(cre)Lowl^*/*J*) were kindly provided by B.B. Lowell (JAX lab stock 016963 and JAX lab stock 016962) [61]; *Som-ires-Cre* mice (*Sst ^tm2.1(cre)Zjh^/J* (JAX lab stock 013044) were kindly donated by Z. J. Huang [62].

Mice were housed at constant temperature (22 ± 1 °C), humidity and circadian cycle (reversed 12 h light-dark cycle). Food and water were available *ad libitum*. Both male and female heterozygous mice 10-12 weeks of age at the start of experimental procedures were used in all experiments.

## Method details

### Viral constructs and preparation

Cre-inducible recombinant AAV vectors carrying transgenes encoding chemogenetic receptors (*pAAV-hSyn-DIO-hM3Dq-mCherry* and *pAAV-hSyn-DIO-hM4Di-mCherry*) were gifts from Bryan L. Roth (Addgene plasmids 44361 and 44362) [63]. For calcium photometry experiments, plasmid *pAAV-hSyn-DIO-GCaMP6s* (Addgene plasmid 184284) was generated by inserting the *GCaMP6s* open reading frame from *pGP-CMV-GCaMP6s* (Addgene plasmid 40753, gift of Douglas Kim) [64] into the *pAAV-hSyn-DIO-hM3Dq-mCherry* backbone, as we described previously [42]. For the caspase lesions, *pAAV-EF1α-DIO-taCASP3-TEV* was a gift from Nirao Shah (Addgene plasmid 45580) [65]. All vectors were packed into AAV capsids in-house (mixed 1:1 ratio of AAV1 and AAV2 capsids proteins with AAV2 ITRs) as described previously [66, 67]. Aliquots of virus were stored at -80 °C before stereotaxic injection.

### Stereotaxic surgery

Mice were anesthetized with isoflurane (induced at 4% and maintained at 1.5% - 2.5%) in oxygen (vol/vol) and positioned on a stereotaxic frame (Angle Two, Leica Microsystems, Milton Keynes, Buckinghamshire, UK). A heat pad was used during the surgery to prevent heat loss (ThermoStar, RWD Life Sciences). A digital mouse brain atlas was linked to the injection frame guide the desired brain regions. The following coordinates were used: for LHb: AP = -1.7 mm; ML = ±0.4 mm; DV = -2.6 mm; for EP: AP = -1.22 mm; ML ±1.77 mm; DV -4.64 mm; for VTA: AP = -3.52 mm; ML -0.36 mm; DV -4.3 mm; for DRN: AP = −4.6 mm; ML = 0.00 mm; DV = −4.64 mm. The virus (0.2 µl – 0.3 µl total volume depending on the viral titer) was injected through a stainless steel 33-gauge/15mm/PST3 internal cannula (Hamilton) attached to a 10 µl Hamilton syringe, at a rate of 0.1 µl min^−1^. After infusion, the cannula was left at the injection site for five minutes and then slowly withdrawn.

For photometry recordings, following AAV injection, a mono-fiber optic cannula (200 µm; Doric Lenses, Inc., Quebec, Canada) was placed 100 μm above the EP, LHb, VTA or DRN. The cannula was affixed with a M1 screw and with dental cement. For locally infusing saline or CNO in the VTA, guide cannulas were implanted above VTA. We conducted post-mortem analysis to verify the viral injection sites and to evaluate the placement of the fiber optic or guide cannulas. Only mice with appropriate viral expression pattern and fibre/cannula implantation were included in this study.

### EEG and EMG recording and scoring of sleep-wake behaviors

Non-tethered EEG and EMG recordings were captured using Neurologger 2A device as described previously [68]. Mice were implanted with three miniature screw electrodes (–1.5 mm Bregma, +1.5 mm midline, first recording electrode; +1.5 mm Bregma, –1.5 mm midline, second recording electrode; –1 mm Lambda, 0 mm midline, reference electrode) with two EMG wires inserted in the dorsal neck muscles (AS634, Cooner Wire, CA). The EEG-EMG device was affixed to the skull with Orthodontic Resin power and Orthodontic resin liquid (Tocdental, UK). All mice were allowed 3-4 weeks for recovery in their home cages. Data were recorded with four times oversampling at a sampling rate of 200 Hz. Collected data was downloaded and waveforms visualized using Spike2 software (Cambridge Electronic Design, Cambridge, UK). The EEG signals were high-pass filtered (0.5 Hz, -3dB, an FFT size of 512 was the designated time window) using a digital filter. The EMG signals were band-pass filtered between 5-45 Hz (-3dB). Power in the delta (0.5-4 Hz), theta (6-10 Hz) bands and theta to delta band ratio were calculated, along with the root mean square (RMS) value of the EMG signal (averaged over a bin size of 5 s). All of these data were used to define the vigilance states of WAKE, NREM and REM by an automatic script. Each vigilance state was screened and confirmed manually afterward. The peak frequency during NREM epochs were analyzed using Fourier transform power spectra to average power spectra over blocks of time.

For chemogenetic experiments [69], clozapine-N-oxide (C0832, Sigma-Aldrich, dissolved in saline, 1 mg/kg) or saline was injected *i.p*. and the vigilance states were recorded. Mice were split into random groups that received either saline or CNO injection. After conducting preliminary experiments, for the hM3-expressing mice, CNO or saline were injected during the “lights off” active phase; for the hM4-expressing mice, CNO or saline were injected at the start of “lights on” sleep phase.

For local chemogenetic activation experiments, CNO (3 µM, 0.2 µl) or saline (0.2 µl) was unilaterally infused via guided cannula over 5min and sleep-wake states were recorded simultaneously.

### Immunohistochemistry

Mice were deeply anesthetized with an overdose of pentobarbital (100 mg/kg body weight; *i.p.*) and transcardially perfused with 4% paraformaldehyde (Thermo scientific) in phosphate buffered saline (PBS, pH 7.4, Sigma-Aldrich). After post-fixation overnight, the mouse brain was preserved in 30% sucrose/PBS for 2 days. 60-μm-thick coronal sections were sliced using a Leica SM 2010R microtome. Free-floating sections were washed in PBS three times for 5 min, permeabilized in PBS plus 0.4% Triton X-100 for 15 min and blocked by incubation in PBS plus 5% normal goat serum (NGS) (Vector), 0.2% Triton X-100 for 1 hour. Sections were incubated with primary antibody diluted in PBS plus 2% NGS overnight at 4°C in a shaker. Incubated slices were then washed three times in PBS for 10 min, and incubated for 2 hours with secondary antibody (Molecular Probes) in PBS and subsequently washed three times in PBS for 10 min (all at room temperature). Before mounting, slices were incubated with Hoechst 33342 (Thermo Fisher Scientific) for 15min followed by rinsing with PBS. Finally, slices were mounted on slides, embedded in Dako mounting medium (Agilent Technologies) and imaged using an inverted wildfield microscope (Zeiss Axio Obseever) or a Leica SP5 MP confocal microscope (Facility for Imaging by Light Microscopy, FILM, Imperial College London).

Primary antibodies used were rabbit polyclonal c-Fos (1:4000, Sigma-Aldrich); rat monoclonal mCherry (1:2000, Thermo Fisher Scientific); chicken polyclonal GFP (1:1000, Abcam); Secondary antibodies were Alexa Fluor 488 goat anti-rabbit, Alexa Fluor 488 goat anti-mouse, Alexa Fluor 594 goat anti-rabbit, Alexa Fluor 594 goat anti-mouse (1:1000, Invitrogen Molecular Probes, UK).

### Fiber photometry

To determine the spontaneous activity of targeted neurons during baseline sleep, we used fiber photometry as previously described [42, 70]. Briefly, we used a 473-nm diode-pumped solid-state blue laser for GCaMP6s excitation (Shanghai Laser & Optics Century Co.). The laser light was passed through a single-source fluorescence cube (FMC_GFP_FC, Doric Lenses) through an optical fiber patch cord (Ø 200 µm, 0.22 numerical aperture, Doric Lenses). From the filter cube, a multimodal optical patch cord (Ø 200 µm, 0.37 numerical aperture) was attached to a ceramic optical fiber (Ø 200 µm, 0.37 numerical aperture) implanted into the mouse brain with a ceramic split mating sleeve ferrule (Thorlabs). The GCaMP6 output was then filtered at 500–550 nm using a second dichroic in the fluorescence cube and converted to voltage by an amplified photodiode (APD-FC, Doric Lenses). The photodiode signal was output to a lock-in amplifier (SR810, Stanford Research Systems) and the power of the laser was set to 80 µW at the fiber tip. The signal was then digitized using a CED 1401 Micro Box (Cambridge Electronic Design) and recorded at 1 kHz using Spike2 software (Cambridge Electronic Design).

The photometry signal was aligned with the EEG and EMG recordings. For each experiment, the photometry signal *F* was normalized to the baseline signal using Δ*F*/*F*(*t*) = (*F*(*t*) − median (*F*))/median (*F*) [42]. We observed a decay of photometry signal at the beginning of some recordings. All the sessions were selected after the photometry signal became stable. We performed the recordings in 2–3 sessions per mouse, one session for 6 h. For the transitions for vigilance states, several sessions were randomly chosen and analyzed.

### Behavioral Tests

To evaluate the behavioral changes after 4-day REM sleep restriction, we used the several assays. All mice were handled daily by the experimenter for 5 minutes for 3 days before the start of the experiments. One the day of testing, mice were transferred to the testing room 30 min prior to the experiment. All behavior tests were performed during the dark phase.

#### Light/dark box assay

The arena (36 (l) x 27 (w) x 30 cm (h), Zantiks LT system) is divided into one dark “safe” compartment (one third) and one large illuminated light compartment (two thirds) with a 50 mm semi-circle hole allowing mice to move freely between the two compartments. Mice were placed in the dark side of the arena and movement was tracked for 5 min by an overhead video camera positioned above the arena. The number of transitions as well as the time spent in each compartment were calculated.

#### Elevated plus maze assay

This was performed as we previously described [71]. The elevated plus maze (Ugo Basile) was made of grey non-reflective plastic and consisted of two open arms (35 x 5 cm) and two enclosed arms (35 x 5 x 15 cm) extending from a central platform at 90 degrees in the form of a plus. The maze was elevated 60 cm from the floor. Individual mice were placed in the center and allowed 5 minutes to explore the maze. The movements were captured and tracked with an overhead video camera. Time spent in each arm, speed and travel distance were calculated using Any-maze software (Stoelting).

#### Looming test

We adopted a previously reported protocol [45]. Briefly, the test was performed in a 48 x 35 x 30 cm arena. The loom stimulus was a 2.5 cm black disk generated using Raspberry Pi. The stimuli were presented on one side of an LCD monitor displaying a grey screen and smoothly moved to the opposite side over 4s. Videos of mouse movements were recorded and processed in MATLAB with a home-made script.

#### Marble burying test

New cages with equal amount of bedding were filled with 20 glass marbles (15-mm diameter) in a 4 x 5 arrangement. Mice were transferred from home cages and allowed to explore the new cages for 20 min. The number of marbles buried (> 2/3 covered by bedding material) was recorded and compared [72].

### Statistical analysis

Mice with an incorrect viral expression or fiber/cannula placement were excluded from the analysis. Analyses were performed with GraphPad Prism 8 (GraphPad software) and MATLAB (Mathworks). Data are presented as the mean ± SEM. We used either two-tailed or paired *t*-test for data that were not independent. *P*-values are shown when they were <0.05.

### Data and code availability

Data reported in this study and any additional information is available from the corresponding authors upon request. This paper does not report original code.

## ACKNOWLEDGMENTS

This research supported by the Wellcome Trust (107839/Z/15/Z, 107841/Z/15/Z and 220759/Z/20/Z, N.P.F. and W.W.), the UK Dementia Research Institute grant UK DRI-5004 (W.W., N.P.F.), the Rubicon Research Program of the Netherlands Organization for Scientific Research (NWO) (019.161LW.010, W.B.) and the People Program (Marie Curie Actions) of the European Union’s Eight Framework Program H2020 under REA grant agreement 753548 (W.B.). The Facility for Imaging by Light Microscopy (FILM) at Imperial College London was in part supported by funding from the Wellcome Trust (grant 104931/Z/14/Z) and BBSRC (grant BB/L015129/1). We thank A. Bruckbauer for the technical support with the microscopes; V. Navetat for helping with some of the illustrations; M. Stephenson-Jones, F. Marbach and L. Rollik (Sainsbury Wellcome Center, UCL) for valuable discussions. For the purpose of open access, the author has applied a CC BY public copyright licence to any Author Accepted Manuscript version arising from this submission.

## AUTHOR CONTRIBUTIONS

W.B., W.W. and N.P.F wrote the manuscript; W.B. designed and performed most experiments and data analysis; M.N., X.Y. and Y.M. performed some behavioral tests; E.B. and E.C.H. performed some data analysis; S.W., A.M. and R.Y. performed some histology experiments; A.L.V developed the neurologgers.

## DECLARATION OF INTERESTS

The authors declare no competing interests.

## Notes

### Competing Interest Statement

The authors have declared no competing interest.

## References

1. Franks, N.P., and Wisden, W. (2021). The inescapable drive to sleep: Overlapping mechanisms of sleep and sedation. Science 374, 556–559.

2. Stephan, A.M., Lecci, S., Cataldi, J., and Siclari, F. (2021). Conscious experiences and high-density EEG patterns predicting subjective sleep depth. Curr Biol 31, 5487–5500 e5483.

3. Della Monica, C., Johnsen, S., Atzori, G., Groeger, J.A., and Dijk, D.J. (2018). Rapid Eye Movement Sleep, Sleep Continuity and Slow Wave Sleep as Predictors of Cognition, Mood, and Subjective Sleep Quality in Healthy Men and Women, Aged 20–84 Years. Front Psychiatry 9, 255.

4. Tsai, C.J., Nagata, T., Liu, C.Y., Suganuma, T., Kanda, T., Miyazaki, T., Liu, K., Saitoh, T., Nagase, H., Lazarus, M., et al. (2021). Cerebral capillary blood flow upsurge during REM sleep is mediated by A2a receptors. Cell Rep 36, 109558.

5. Nollet, M., Hicks, H., McCarthy, A.P., Wu, H., Moller-Levet, C.S., Laing, E.E., Malki, K., Lawless, N., Wafford, K.A., Dijk, D.J., et al. (2019). REM sleep’s unique associations with corticosterone regulation, apoptotic pathways, and behavior in chronic stress in mice. Proc Natl Acad Sci U S A 116, 2733–2742.

6. Wassing, R., Benjamins, J.S., Dekker, K., Moens, S., Spiegelhalder, K., Feige, B., Riemann, D., van der Sluis, S., Van Der Werf, Y.D., Talamini, L.M., et al. (2016). Slow dissolving of emotional distress contributes to hyperarousal. Proc Natl Acad Sci U S A 113, 2538–2543.

7. Goldstein, A.N., and Walker, M.P. (2014). The role of sleep in emotional brain function. Annu Rev Clin Psychol 10, 679–708.

8. Hutchison, I.C., and Rathore, S. (2015). The role of REM sleep theta activity in emotional memory. Front Psychol 6, 1439.

9. Pace-Schott, E.F., Germain, A., and Milad, M.R. (2015). Effects of sleep on memory for conditioned fear and fear extinction. Psychol Bull 141, 835–857.

10. Wassing, R., Lakbila-Kamal, O., Ramautar, J.R., Stoffers, D., Schalkwijk, F., and Van Someren, E.J.W. (2019). Restless REM Sleep Impedes Overnight Amygdala Adaptation. Curr Biol 29, 2351–2358 e2354.

11. Miller, K.E., and Gehrman, P.R. (2019). REM Sleep: What Is It Good For? Curr Biol 29, R806–R807.

12. Murkar, A.L.A., and De Koninck, J. (2018). Consolidative mechanisms of emotional processing in REM sleep and PTSD. Sleep Med Rev 41, 173–184.

13. Simionato, N.M., da Silva Rocha-Lopes, J., Machado, R.B., and Suchecki, D. (2022). Chronic rapid eye movement sleep restriction during juvenility has long-term effects on anxiety-like behaviour and neurotransmission of male Wistar rats. Pharmacol Biochem Behav 217, 173410.

14. Boyce, R., Glasgow, S.D., Williams, S., and Adamantidis, A. (2016). Causal evidence for the role of REM sleep theta rhythm in contextual memory consolidation. Science 352, 812–816.

15. Aime, M., Calcini, N., Borsa, M., Campelo, T., Rusterholz, T., Sattin, A., Fellin, T., and Adamantidis, A. (2022). Paradoxical somatodendritic decoupling supports cortical plasticity during REM sleep. Science 376, 724–730.

16. Izawa, S., Chowdhury, S., Miyazaki, T., Mukai, Y., Ono, D., Inoue, R., Ohmura, Y., Mizoguchi, H., Kimura, K., Yoshioka, M., et al. (2019). REM sleep-active MCH neurons are involved in forgetting hippocampus-dependent memories. Science 365, 1308–1313.

17. Tseng, Y.T., Zhao, B., Chen, S., Ye, J., Liu, J., Liang, L., Ding, H., Schaefke, B., Yang, Q., Wang, L., et al. (2022). The subthalamic corticotropin-releasing hormone neurons mediate adaptive REM-sleep responses to threat. Neuron 110, 1223–1239 e1228.

18. Palagini, L., Baglioni, C., Ciapparelli, A., Gemignani, A., and Riemann, D. (2013). REM sleep dysregulation in depression: state of the art. Sleep Med Rev 17, 377–390.

19. Kudlow, P.A., Cha, D.S., Lam, R.W., and McIntyre, R.S. (2013). Sleep architecture variation: a mediator of metabolic disturbance in individuals with major depressive disorder. Sleep Med 14, 943–949.

20. Riemann, D., Krone, L.B., Wulff, K., and Nissen, C. (2020). Sleep, insomnia, and depression. Neuropsychopharmacology 45, 74–89.

21. Peever, J., and Fuller, P.M. (2017). The Biology of REM Sleep. Curr Biol 27, R1237–R1248.

22. Chen, K.S., Xu, M., Zhang, Z., Chang, W.C., Gaj, T., Schaffer, D.V., and Dan, Y. (2018). A Hypothalamic Switch for REM and Non-REM Sleep. Neuron 97, 1168–1176 e1164.

23. Luppi, P.H., Billwiller, F., and Fort, P. (2017). Selective activation of a few limbic structures during paradoxical (REM) sleep by the claustrum and the supramammillary nucleus: evidence and function. Curr Opin Neurobiol 44, 59–64.

24. Renouard, L., Billwiller, F., Ogawa, K., Clement, O., Camargo, N., Abdelkarim, M., Gay, N., Scote-Blachon, C., Toure, R., Libourel, P.A., et al. (2015). The supramammillary nucleus and the claustrum activate the cortex during REM sleep. Sci Adv 1, e1400177.

25. Gelegen, C., Miracca, G., Ran, M.Z., Harding, E.C., Ye, Z., Yu, X., Tossell, K., Houston, C.M., Yustos, R., Hawkins, E.D., et al. (2018). Excitatory Pathways from the Lateral Habenula Enable Propofol-Induced Sedation. Curr Biol 28, 580–587 e585.

26. Stephenson-Jones, M., Yu, K., Ahrens, S., Tucciarone, J.M., van Huijstee, A.N., Mejia, L.A., Penzo, M.A., Tai, L.H., Wilbrecht, L., and Li, B. (2016). A basal ganglia circuit for evaluating action outcomes. Nature 539, 289–293.

27. Cerniauskas, I., Winterer, J., de Jong, J.W., Lukacsovich, D., Yang, H., Khan, F., Peck, J.R., Obayashi, S.K., Lilascharoen, V., Lim, B.K., et al. (2019). Chronic Stress Induces Activity, Synaptic, and Transcriptional Remodeling of the Lateral Habenula Associated with Deficits in Motivated Behaviors. Neuron 104, 899–915 e898.

28. Li, H., Pullmann, D., and Jhou, T.C. (2019). Valence-encoding in the lateral habenula arises from the entopeduncular region. Elife 8.

29. Hong, S., and Hikosaka, O. (2008). The globus pallidus sends reward-related signals to the lateral habenula. Neuron 60, 720–729.

30. Hu, H., Cui, Y., and Yang, Y. (2020). Circuits and functions of the lateral habenula in health and in disease. Nat Rev Neurosci 21, 277–295.

31. Hikosaka, O. (2010). The habenula: from stress evasion to value-based decision-making. Nat Rev Neurosci 11, 503–513.

32. Gordon-Fennell, A., and Stuber, G.D. (2021). Illuminating subcortical GABAergic and glutamatergic circuits for reward and aversion. Neuropharmacology 198, 108725.

33. Zhang, L., Hernandez, V.S., Swinny, J.D., Verma, A.K., Giesecke, T., Emery, A.C., Mutig, K., Garcia-Segura, L.M., and Eiden, L.E. (2018). A GABAergic cell type in the lateral habenula links hypothalamic homeostatic and midbrain motivation circuits with sex steroid signaling. Transl Psychiatry 8, 50.

34. Namboodiri, V.M., Rodriguez-Romaguera, J., and Stuber, G.D. (2016). The habenula. Curr Biol 26, R873–R877.

35. Xu, J., Jo, A., DeVries, R.P., Deniz, S., Cherian, S., Sunmola, I., Song, X., Marshall, J.J., Gruner, K.A., Daigle, T.L., et al. (2022). Intersectional mapping of multi-transmitter neurons and other cell types in the brain. Cell Rep 40, 111036.

36. Shabel, S.J., Proulx, C.D., Trias, A., Murphy, R.T., and Malinow, R. (2012). Input to the lateral habenula from the basal ganglia is excitatory, aversive, and suppressed by serotonin. Neuron 74, 475–481.

37. Parent, M., Levesque, M., and Parent, A. (2001). Two types of projection neurons in the internal pallidum of primates: single-axon tracing and three-dimensional reconstruction. J Comp Neurol 439, 162–175.

38. Kim, S., Wallace, M.L., El-Rifai, M., Knudsen, A.R., and Sabatini, B.L. (2022). Co-packaging of opposing neurotransmitters in individual synaptic vesicles in the central nervous system. Neuron 110, 1371–1384 e1377.

39. Wallace, M.L., Saunders, A., Huang, K.W., Philson, A.C., Goldman, M., Macosko, E.Z., McCarroll, S.A., and Sabatini, B.L. (2017). Genetically Distinct Parallel Pathways in the Entopeduncular Nucleus for Limbic and Sensorimotor Output of the Basal Ganglia. Neuron 94, 138–152 e135.

40. Hernandez-Martinez, R., and Calakos, N. (2017). Seq-ing the Circuit Logic of the Basal Ganglia. Trends Neurosci 40, 325–327.

41. Lazaridis, I., Tzortzi, O., Weglage, M., Martin, A., Xuan, Y., Parent, M., Johansson, Y., Fuzik, J., Furth, D., Fenno, L.E., et al. (2019). A hypothalamus-habenula circuit controls aversion. Mol Psychiatry 24, 1351–1368.

42. Yu, X., Li, W., Ma, Y., Tossell, K., Harris, J.J., Harding, E.C., Ba, W., Miracca, G., Wang, D., Li, L., et al. (2019). GABA and glutamate neurons in the VTA regulate sleep and wakefulness. Nat Neurosci 22, 106–119.

43. Oikonomou, G., Altermatt, M., Zhang, R.W., Coughlin, G.M., Montz, C., Gradinaru, V., and Prober, D.A. (2019). The Serotonergic Raphe Promote Sleep in Zebrafish and Mice. Neuron 103, 686–701 e688.

44. Takahashi, A., Durand-de Cuttoli, R., Flanigan, M.E., Hasegawa, E., Tsunematsu, T., Aleyasin, H., Cherasse, Y., Miya, K., Okada, T., Keino-Masu, K., et al. (2022). Lateral habenula glutamatergic neurons projecting to the dorsal raphe nucleus promote aggressive arousal in mice. Nat Commun 13, 4039.

45. De Franceschi, G., Vivattanasarn, T., Saleem, A.B., and Solomon, S.G. (2016). Vision Guides Selection of Freeze or Flight Defense Strategies in Mice. Curr Biol 26, 2150–2154.

46. Luppi, P.H., and Fort, P. (2019). Neuroanatomical and Neurochemical Bases of Vigilance States. Handb Exp Pharmacol 253, 35–58.

47. Miracca, G., Anuncibay-Soto, B., Tossell, K., Yustos, R., Vyssotski, A.L., Franks, N.P., and Wisden, W. (2022). NMDA Receptors in the Lateral Preoptic Hypothalamus Are Essential for Sustaining NREM and REM Sleep. J Neurosci 42, 5389–5409.

48. Park, S.H., and Weber, F. (2020). Neural and Homeostatic Regulation of REM Sleep. Front Psychol 11, 1662.

49. Hasegawa, E., Miyasaka, A., Sakurai, K., Cherasse, Y., Li, Y., and Sakurai, T. (2022). Rapid eye movement sleep is initiated by basolateral amygdala dopamine signaling in mice. Science 375, 994–1000.

50. Oishi, Y., and Lazarus, M. (2017). The control of sleep and wakefulness by mesolimbic dopamine systems. Neurosci Res 118, 66–73.

51. Liu, D., Li, W., Ma, C., Zheng, W., Yao, Y., Tso, C.F., Zhong, P., Chen, X., Song, J.H., Choi, W., et al. (2020). A common hub for sleep and motor control in the substantia nigra. Science 367, 440–445.

52. Qiu, M.H., Zhong, Z.G., Chen, M.C., and Lu, J. (2019). Nigrostriatal and mesolimbic control of sleep-wake behavior in rat. Brain Struct Funct 224, 2525–2535.

53. Qiu, M.H., Chen, M.C., Wu, J., Nelson, D., and Lu, J. (2016). Deep brain stimulation in the globus pallidus externa promotes sleep. Neuroscience 322, 115–120.

54. Proulx, C.D., Hikosaka, O., and Malinow, R. (2014). Reward processing by the lateral habenula in normal and depressive behaviors. Nat Neurosci 17, 1146–1152.

55. Szonyi, A., Zicho, K., Barth, A.M., Gonczi, R.T., Schlingloff, D., Torok, B., Sipos, E., Major, A., Bardoczi, Z., Sos, K.E., et al. (2019). Median raphe controls acquisition of negative experience in the mouse. Science 366.

56. Andalman, A.S., Burns, V.M., Lovett-Barron, M., Broxton, M., Poole, B., Yang, S.J., Grosenick, L., Lerner, T.N., Chen, R., Benster, T., et al. (2019). Neuronal Dynamics Regulating Brain and Behavioral State Transitions. Cell 177, 970–985 e920.

57. Yu, X., Zhao, G., Wang, D., Wang, S., Li, R., Li, A., Wang, H., Nollet, M., Chun, Y.Y., Zhao, T., et al. (2022). A specific circuit in the midbrain detects stress and induces restorative sleep. Science 377, 63–72.

58. Meye, F.J., Soiza-Reilly, M., Smit, T., Diana, M.A., Schwarz, M.K., and Mameli, M. (2016). Shifted pallidal co-release of GABA and glutamate in habenula drives cocaine withdrawal and relapse. Nat Neurosci 19, 1019–1024.

59. Shabel, S.J., Proulx, C.D., Piriz, J., and Malinow, R. (2014). Mood regulation. GABA/glutamate co-release controls habenula output and is modified by antidepressant treatment. Science 345, 1494–1498.

60. Yang, Y., Cui, Y., Sang, K., Dong, Y., Ni, Z., Ma, S., and Hu, H. (2018). Ketamine blocks bursting in the lateral habenula to rapidly relieve depression. Nature 554, 317–322.

61. Vong, L., Ye, C., Yang, Z., Choi, B., Chua, S., Jr., and Lowell, B.B. (2011). Leptin action on GABAergic neurons prevents obesity and reduces inhibitory tone to POMC neurons. Neuron 71, 142–154.

62. Taniguchi, H., He, M., Wu, P., Kim, S., Paik, R., Sugino, K., Kvitsiani, D., Fu, Y., Lu, J., Lin, Y., et al. (2011). A resource of Cre driver lines for genetic targeting of GABAergic neurons in cerebral cortex. Neuron 71, 995–1013.

63. Krashes, M.J., Koda, S., Ye, C., Rogan, S.C., Adams, A.C., Cusher, D.S., Maratos-Flier, E., Roth, B.L., and Lowell, B.B. (2011). Rapid, reversible activation of AgRP neurons drives feeding behavior in mice. J Clin Invest 121, 1424–1428.

64. Chen, T.W., Wardill, T.J., Sun, Y., Pulver, S.R., Renninger, S.L., Baohan, A., Schreiter, E.R., Kerr, R.A., Orger, M.B., Jayaraman, V., et al. (2013). Ultrasensitive fluorescent proteins for imaging neuronal activity. Nature 499, 295–300.

65. Yang, C.F., Chiang, M.C., Gray, D.C., Prabhakaran, M., Alvarado, M., Juntti, S.A., Unger, E.K., Wells, J.A., and Shah, N.M. (2013). Sexually dimorphic neurons in the ventromedial hypothalamus govern mating in both sexes and aggression in males. Cell 153, 896–909.

66. Murray, A.J., Sauer, J.F., Riedel, G., McClure, C., Ansel, L., Cheyne, L., Bartos, M., Wisden, W., and Wulff, P. (2011). Parvalbumin-positive CA1 interneurons are required for spatial working but not for reference memory. Nat Neurosci 14, 297–299.

67. Klugmann, M., Symes, C.W., Leichtlein, C.B., Klaussner, B.K., Dunning, J., Fong, D., Young, D., and During, M.J. (2005). AAV-mediated hippocampal expression of short and long Homer 1 proteins differentially affect cognition and seizure activity in adult rats. Mol Cell Neurosci 28, 347–360.

68. Anisimov, V.N., Herbst, J.A., Abramchuk, A.N., Latanov, A.V., Hahnloser, R.H., and Vyssotski, A.L. (2014). Reconstruction of vocal interactions in a group of small songbirds. Nat Methods 11, 1135–1137.

69. Alexander, G.M., Rogan, S.C., Abbas, A.I., Armbruster, B.N., Pei, Y., Allen, J.A., Nonneman, R.J., Hartmann, J., Moy, S.S., Nicolelis, M.A., et al. (2009). Remote control of neuronal activity in transgenic mice expressing evolved G protein-coupled receptors. Neuron 63, 27–39.

70. Gunaydin, L.A., Grosenick, L., Finkelstein, J.C., Kauvar, I.V., Fenno, L.E., Adhikari, A., Lammel, S., Mirzabekov, J.J., Airan, R.D., Zalocusky, K.A., et al. (2014). Natural neural projection dynamics underlying social behavior. Cell 157, 1535–1551.

71. Yu, X., Ba, W., Zhao, G., Ma, Y., Harding, E.C., Yin, L., Wang, D., Li, H., Zhang, P., Shi, Y., et al. (2021). Dysfunction of ventral tegmental area GABA neurons causes mania-like behavior. Mol Psychiatry 26, 5213–5228.

72. Thomas, A., Burant, A., Bui, N., Graham, D., Yuva-Paylor, L.A., and Paylor, R. (2009). Marble burying reflects a repetitive and perseverative behavior more than novelty-induced anxiety. Psychopharmacology (Berl) 204, 361–373.

