## Supplemental Figures and legends for "A REM-active basal ganglia circuit that regulates anxiety"

**This PDF file includes:**

Figures S1 to S10

**Figure S1 Histological verification and controls for EP photometry recordings, Related to Figure 1.**

(A) Confirmation of viral injection and optic fiber implantation. Green, areas infected with *AAV-DIO-GCaMP6s*; Blue, location of optic fiber placement. (B) Schematic of the photometry recordings on *Som-Cre* mice injected with *AAV-DIO-GFP* control and representative traces aligned with EEG/EMG recordings. (C) Quantification of the $Ca^{2+}$ signal in each vigilant state showed no vigilance state dependent activity.

**Figure S2 Verification of viral expression of hM4Di and Caspase mediated ablation, Related to Figure 2.**

(A) Confirmation of the expression of *AAV-DIO-hM4Di-mCherry* in the EP. (B) Assessment of $EP^{Som}$ lesioning. Bilateral injection of *AAV-DIO-mCherry* only (control) or combined *AAV-DIO-mCherry* and *AAV-DIO-CASP3* into the EP of *Som-Cre* mice. Representative images show mCherry expression. The fluorescence signal was diminished in the $EP^{Som\text{-}CASP3}$ group ($P = 0.009$, $n = 3$).

**Figure S3 Verification of viral expression and activation of hM3Dq, Related to Figure 2.**

(A) Histological confirmation of *AAV-DIO-hM3Dq-mCherry*. (B) Representative images showing increased cfos expression after CNO injection in the EP. Scale bar, 100 μm.

**Figure S4 $EP^{Som}$ terminals in the LHb are active during REM, mirroring activity in the EP soma, Related to Figure 3.**

*AAV-DIO-GCaMP6s* was injected in $EP^{Som}$ neurons of *Som-Cre* mice and an optical fiber positioned over the LHb. The $Ca^{2+}$ transients at EP→LHb terminals peaked during REM sleep. NREM-REM transitions were associated with prominent increases of activity at these terminals.

**Figure S5 LPO terminals in the LHb are selectively WAKE active, Related to Figure 3.**

*AAV-DIO-GCaMP6s* was injected in $LPO^{Vglut2}$ neurons of *Vglut2-Cre* mice and an optical fiber positioned over the LHb. The $Ca^{2+}$ transients at EP→LHb terminals were highest during wakefulness.

**Figure S6 Changes in the number and duration of REM episodes during activation and inhibition of $LHb^{Vglut2}$ neurons, Related to Figure 4.**

(A) *AAV-DIO-hM4Di-mCherry* was bilaterally injected into the LHb of *Vglut2-Cre* mice. CNO-mediated inhibition of $LHb^{Vglut2}$ neurons decreased REM sleep by decreasing the number of REM episodes. (B) *AAV-DIO-hM3Dq-mCherry* was bilaterally injected into the LHb of *Vglut2-Cre* mice. CNO-mediated excitation of $LHb^{Vglut2}$ neurons increased REM sleep by increasing the number of REM episodes and increasing their durations. The duration of NREM episodes also increased. In both groups of mice, CNO was injected *i.p.* at 1 mg/kg.

**Figure S7 CNO (1 mg/kg; *i.p.*) had no effect on vigilance states in *Vglut2-Cre* mice, Related to Figure 4.**

Following *i.p.* injection, CNO at 1 mg/kg at had no effect on the amounts of WAKE, NREM or REM sleep over the following 24 hours compared with mice injected with saline ($n$ = 6).

**Figure S8 Representative traces of $Ca^{2+}$ signals in the LHb terminals in the dorsal raphe aligned with EEG and EMG recordings during wakefulness, NREM and REM sleep showing increased activity during WAKE, Related to Figure 5.**

Fiber photometry recordings at LHb→DRN terminals across sleep-wake cycle were achieved by expressing *AAV-DIO-GCaMP6s* in $LHb^{Vglut2}$ neurons and with optical fibers placed over terminals in the dorsal raphe (DR).

**Figure S9 Local chemogenetic activation of $\mathbf{LHb^{Vglut2}}$ terminals in the VTA neurons increases REM and NREM sleep, Related to Figure 5.**

(A) Excitation of $LHb^{Vglut2}$ terminals in VTA neurons using hM3Dq-mCherry receptors. *AAV-DIO-hM3Dq-mCherry* was injected into the LHb of *Vglut2-Cre* mice. Immunostaining confirmed the expression of hM3Dq-mCherry in the LHb soma and terminals in the VTA. (Scale bars, 100 μm.) (B) Representative EEG/EMG recordings post-saline or CNO local infusion into the VTA. (C) Summarized changes on WAKE, NREM and REM duration. Data are means ± SEM ($n$ = 7).

**Figure S10 Repeated inhibition of lateral habenula $\mathbf{LHb^{Vglut2}}$ neurons by CNO caused a reduction in REM, Related to Figure 6.**

*Vglut2-Cre* mice were injected bilaterally into the lateral habenula (LHb) with *AAV-DIO-hM4Di-mCherry*. Prior to behavioral assays, the percentage of REM was reduced by repeated (once per day for 4 days) injections of CNO (1 mg/kg) in $LHb^{Vglut2}$ mice.

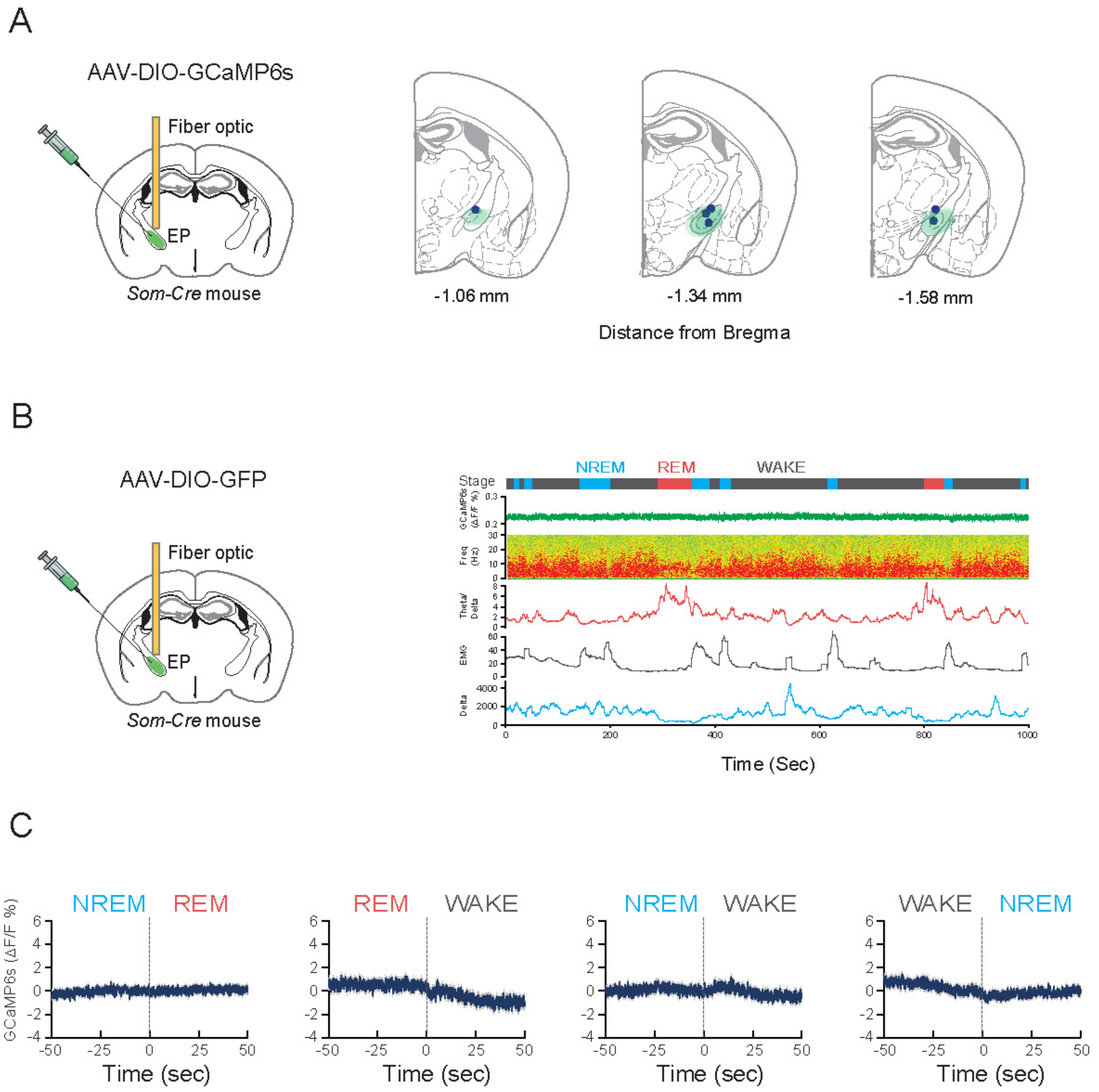
A
AAV-DIO-GCaMP6s
Fiber optic
EP
Som-Cre mouse
-1.06 mm
-1.34 mm
-1.58 mm
Distance from Bregma
B
AAV-DIO-GFP
Fiber optic
EP
Som-Cre mouse
NREM
REM
WAKE
Stage
Time (Sec)
C
NREM
REM
WAKE
GCaMP6s (ΔF/F %)
Time (sec)

Figure S1

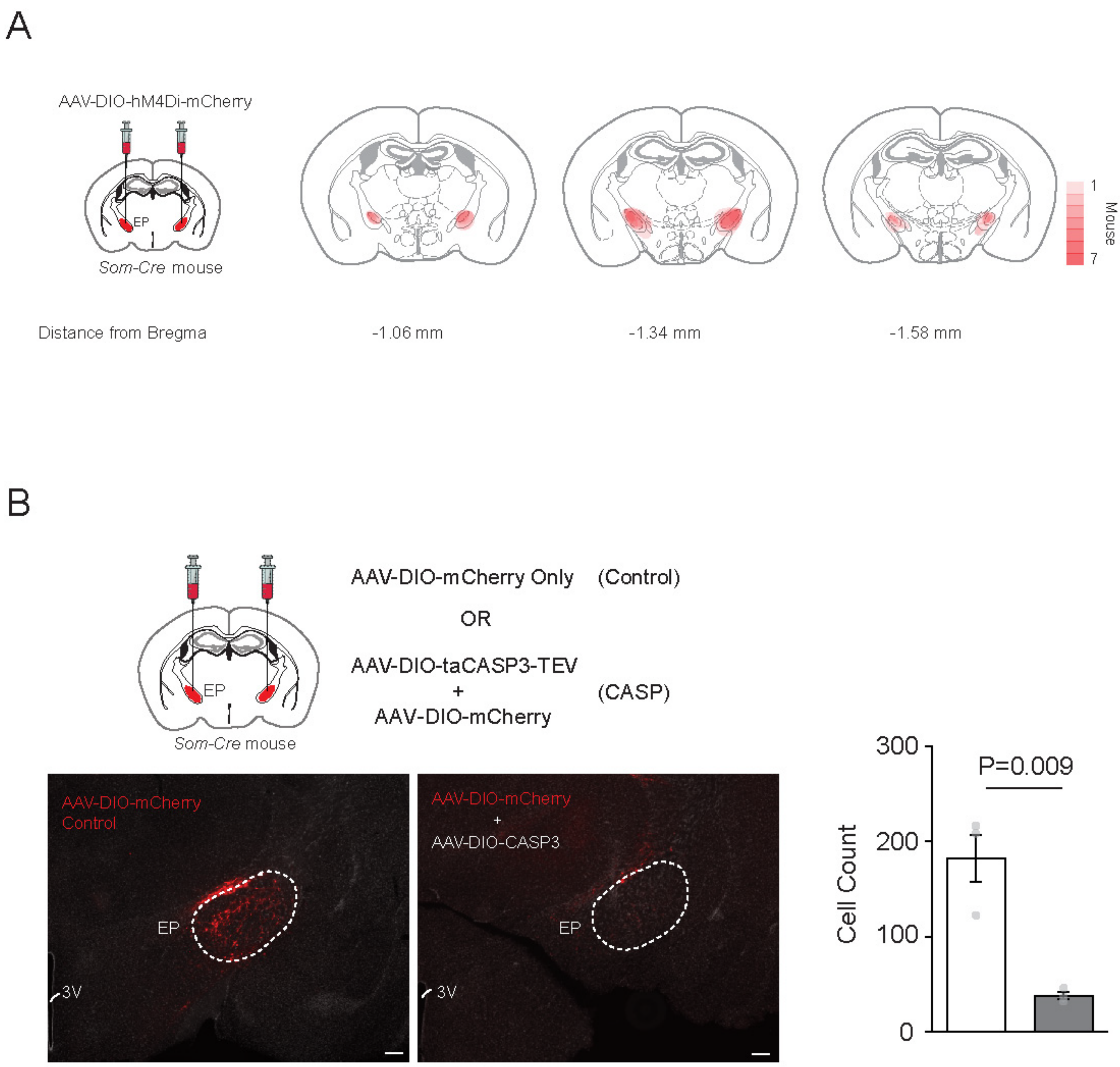
A
AAV-DIO-hM4Di-mCherry
EP
Som-Cre mouse
1
7
Mouse
Distance from Bregma
-1.06 mm
-1.34 mm
-1.58 mm
B
AAV-DIO-mCherry Only (Control)
OR
AAV-DIO-taCASP3-TEV
+
AAV-DIO-mCherry
(CASP)
EP
Som-Cre mouse
AAV-DIO-mCherry
Control
EP
3V
AAV-DIO-mCherry
+
AAV-DIO-CASP3
EP
3V
Cell Count
300
200
100
0
P=0.009

Figure S2

A

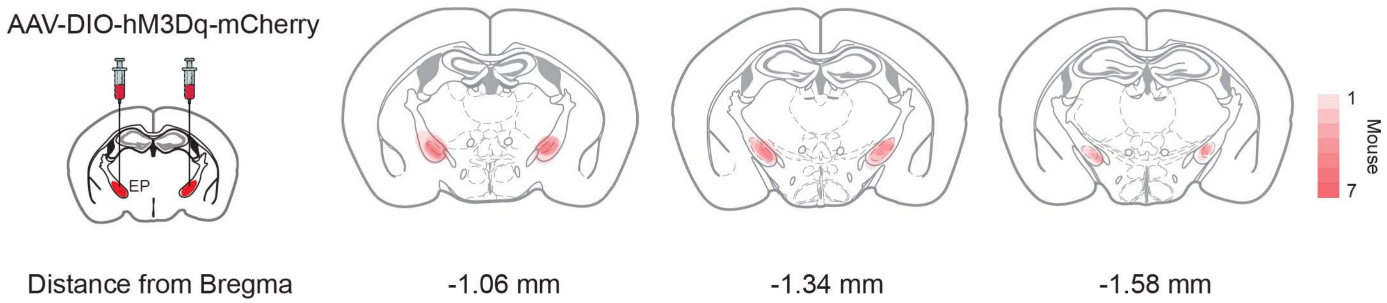
AAV-DIO-hM3Dq-mCherry
EP
1
Mouse
7
Distance from Bregma
-1.06 mm
-1.34 mm
-1.58 mm

B

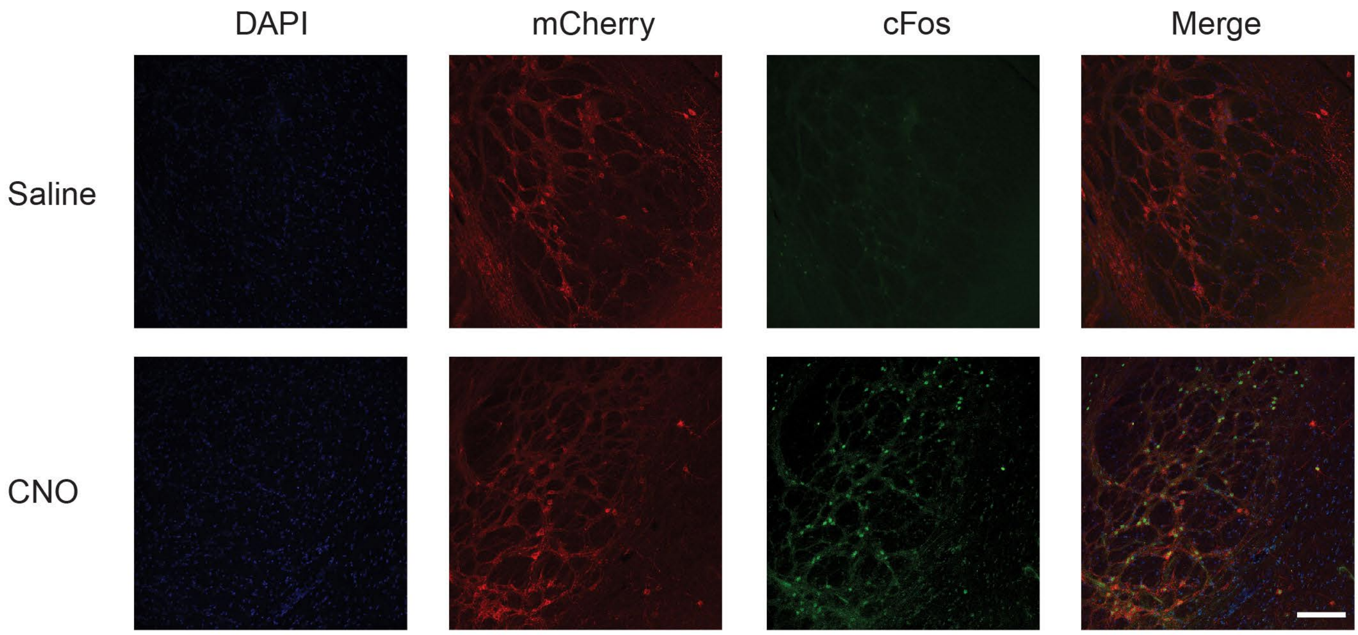
DAPI
mCherry
cFos
Merge
Saline
CNO

Figure S3

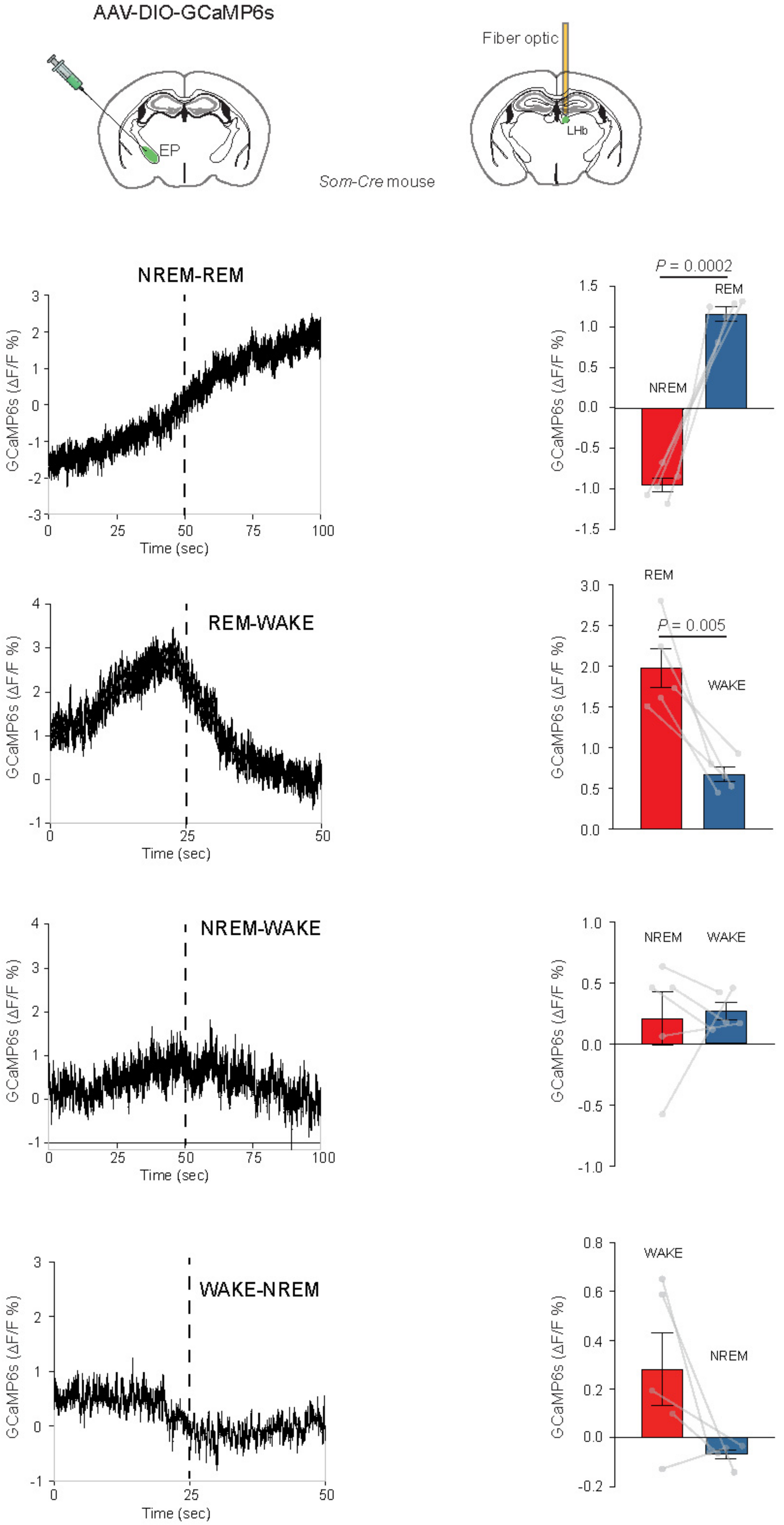
AAV-DIO-GCaMP6s
Fiber optic
EP
LHb
Som-Cre mouse
NREM-REM
REM-WAKE
NREM-WAKE
WAKE-NREM
GCaMP6s (ΔF/F %)
Time (sec)
P = 0.0002
P = 0.005
NREM
REM
WAKE

Figure S4

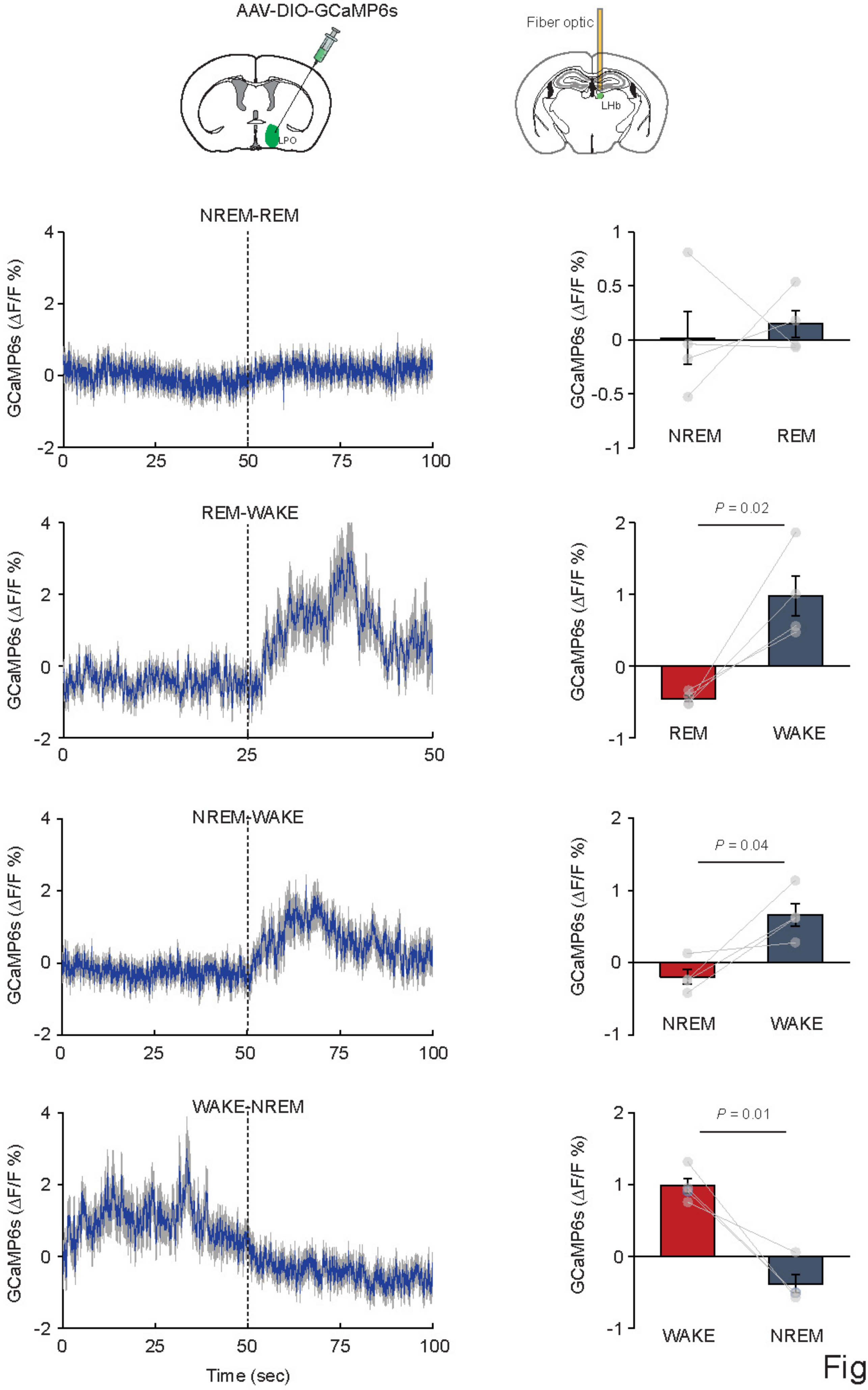
AAV-DIO-GCaMP6s
LPO
Fiber optic
LHb
NREM-REM
REM-WAKE
NREM-WAKE
WAKE-NREM
GCaMP6s (ΔF/F %)
Time (sec)
NREM
REM
WAKE
P = 0.02
P = 0.04
P = 0.01

Figure S5

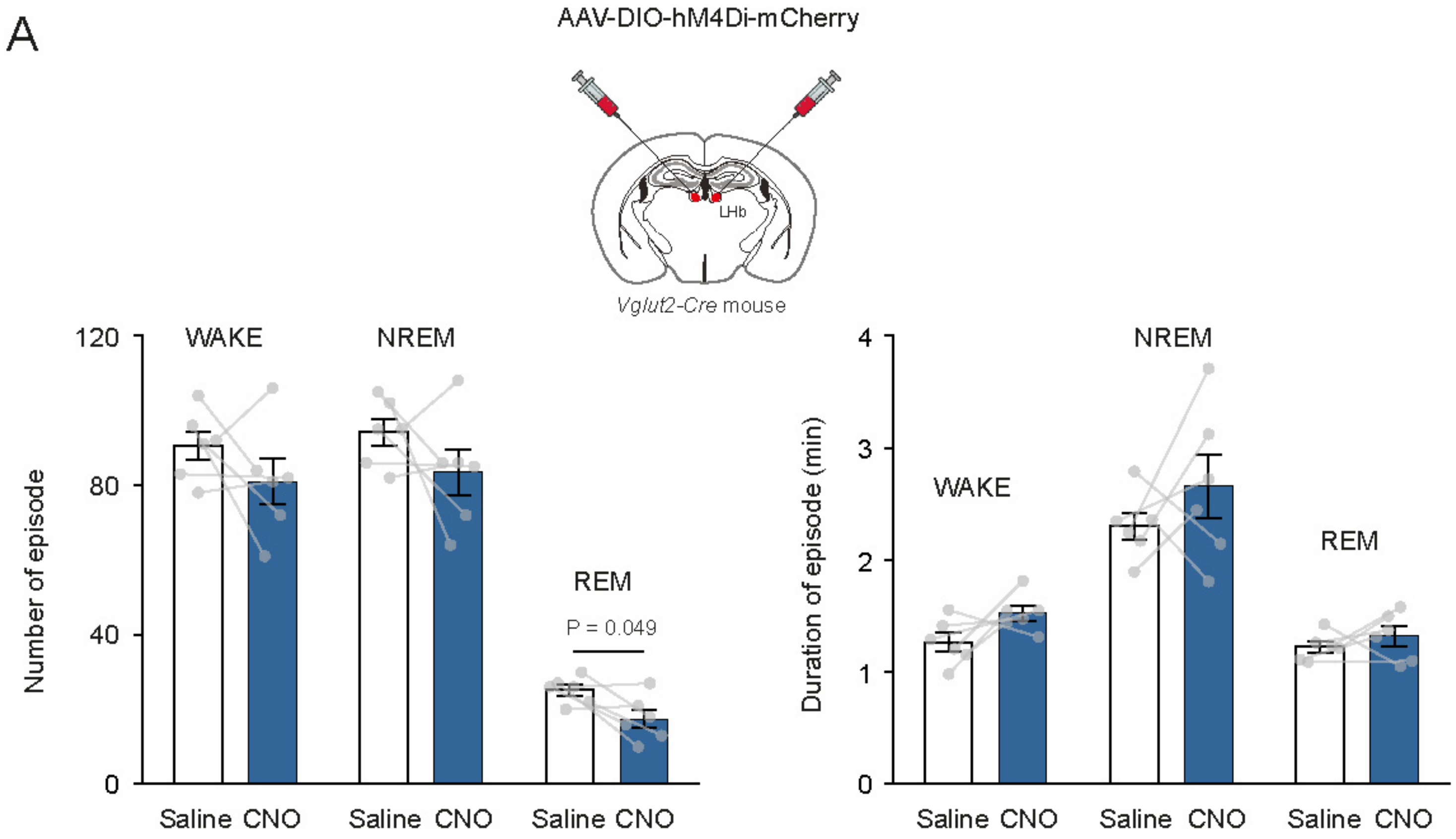
A
AAV-DIO-hM4Di-mCherry
LHb
Vglut2-Cre mouse
Number of episode
120
80
40
0
WAKE
NREM
REM
P = 0.049
Saline CNO
Saline CNO
Saline CNO
Duration of episode (min)
4
3
2
1
0
WAKE
NREM
REM
Saline CNO
Saline CNO
Saline CNO

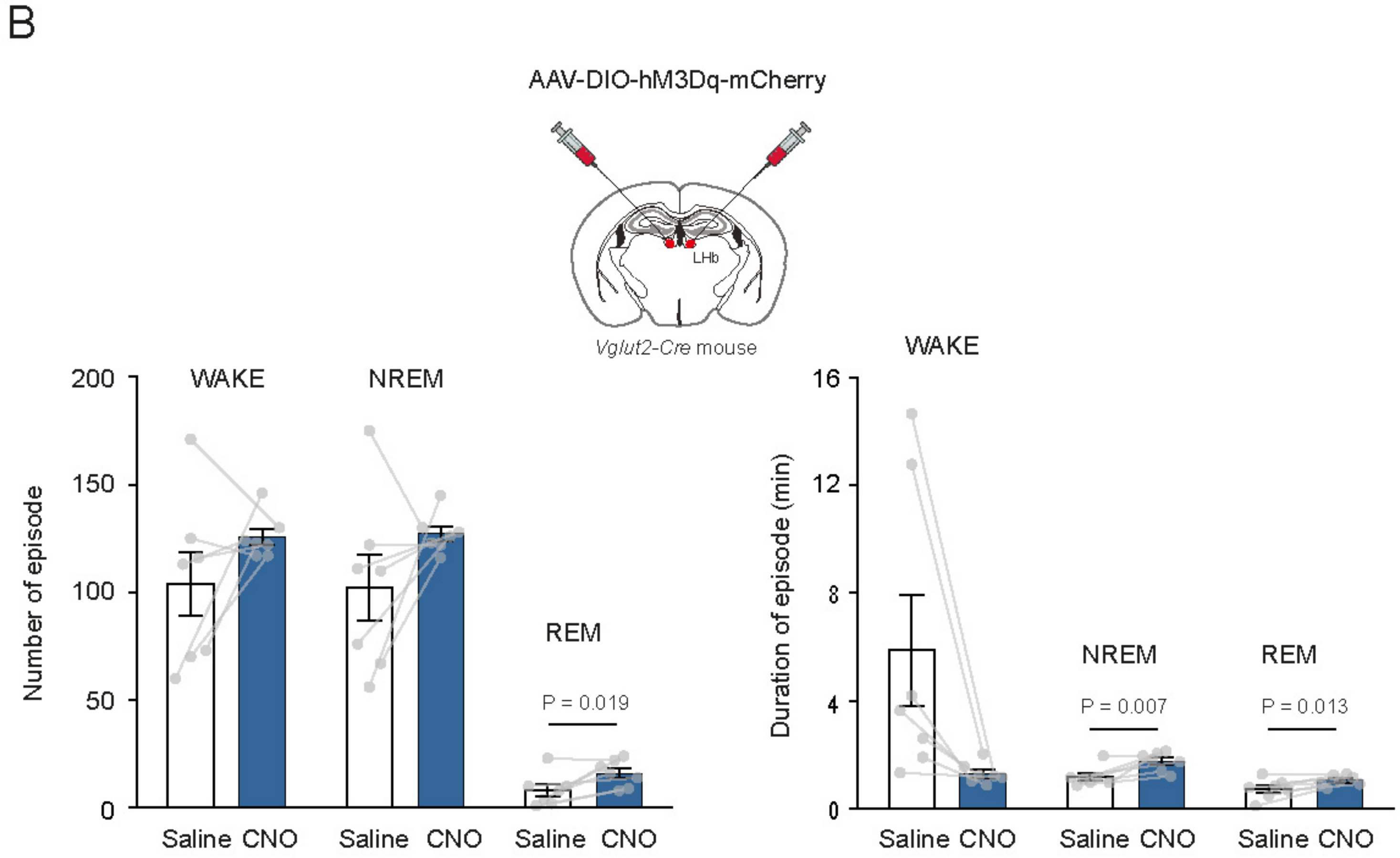
B
AAV-DIO-hM3Dq-mCherry
LHb
Vglut2-Cre mouse
Number of episode
200
150
100
50
0
WAKE
NREM
REM
P = 0.019
Saline CNO
Saline CNO
Saline CNO
Duration of episode (min)
16
12
8
4
0
WAKE
NREM
REM
P = 0.007
P = 0.013
Saline CNO
Saline CNO
Saline CNO

Figure S6

### *Vglut2-Cre* mice

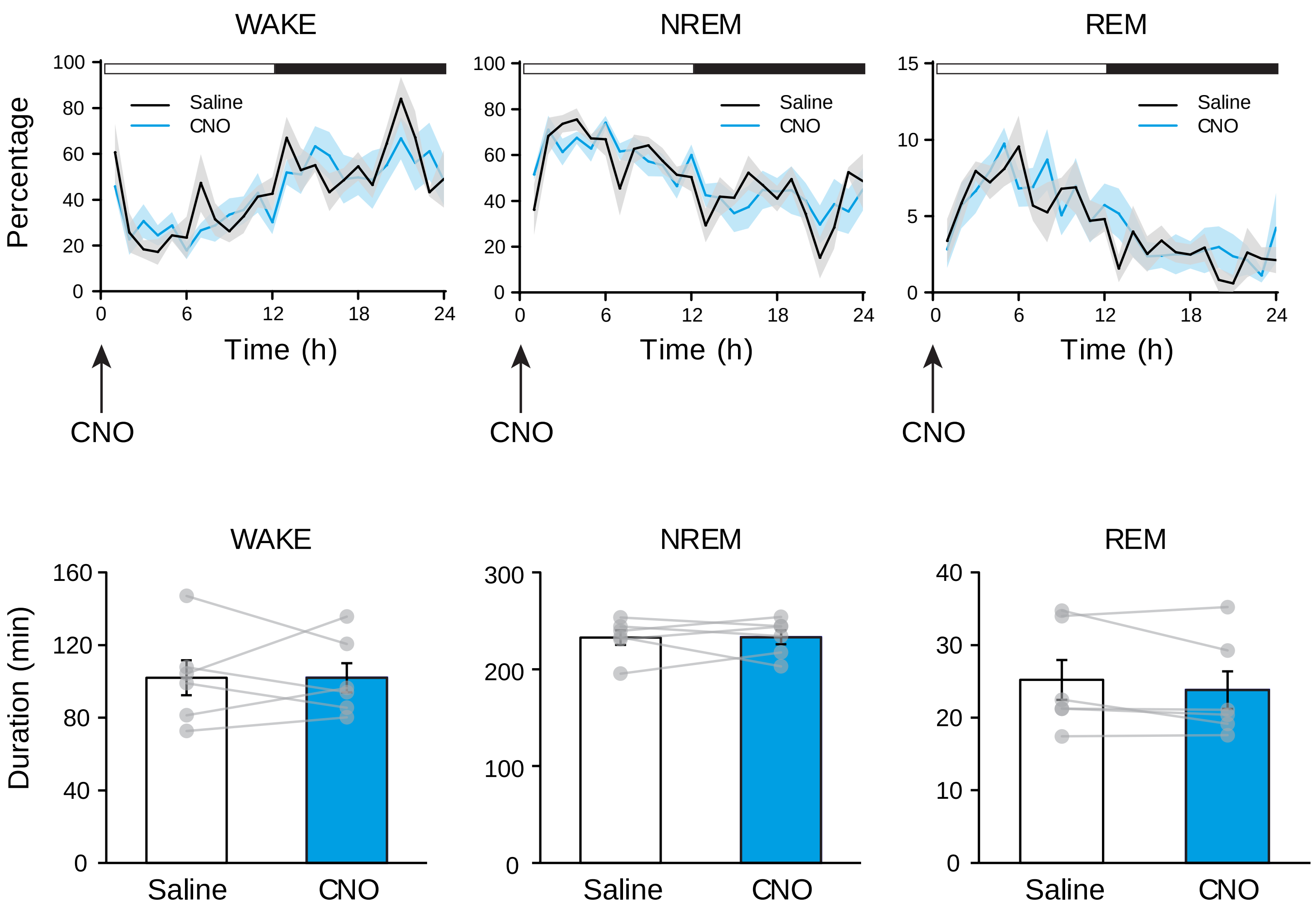


Figure S7

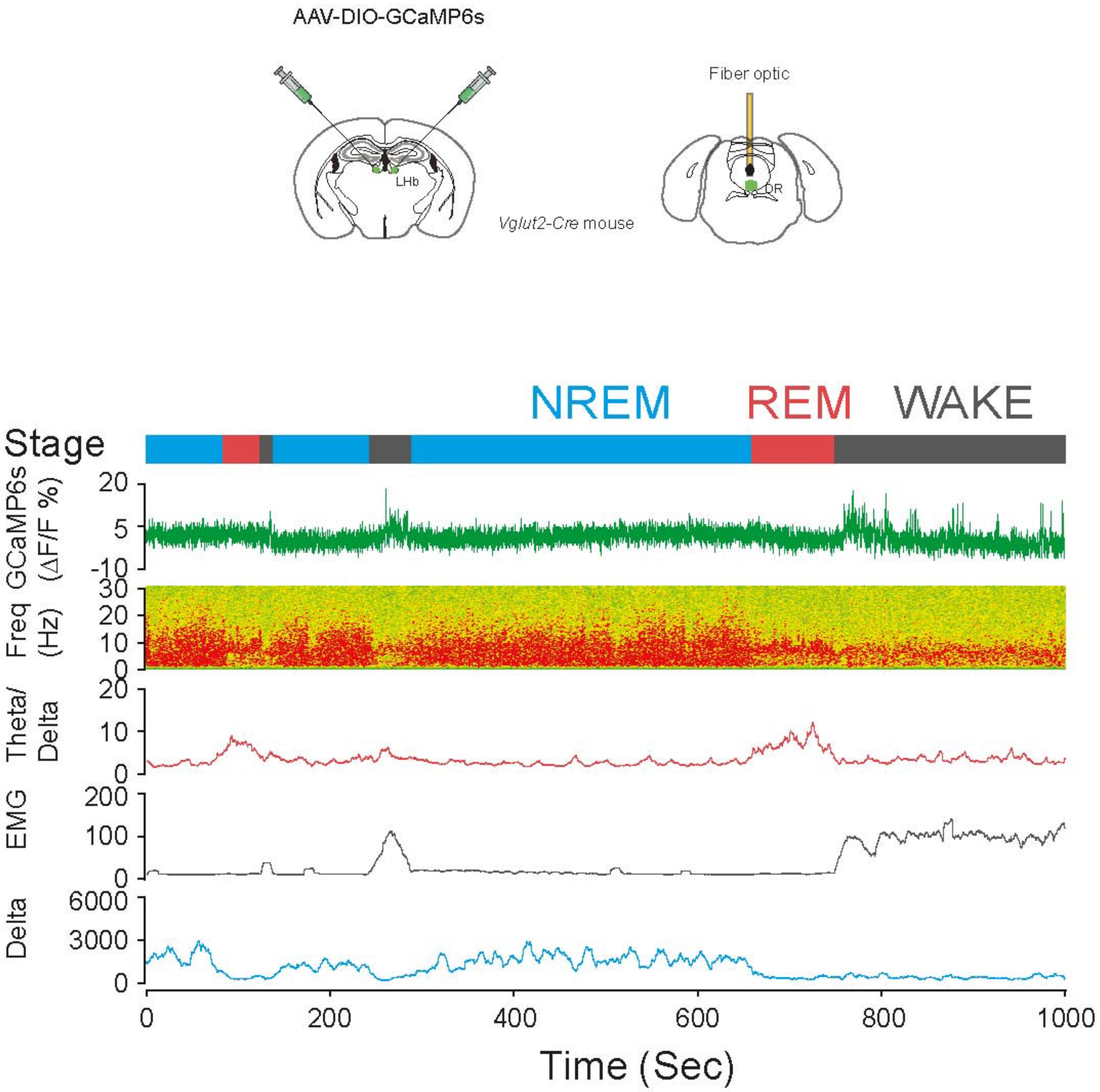
AAV-DIO-GCaMP6s
LHb
Vglut2-Cre mouse
Fiber optic
DR
NREM
REM
WAKE
Stage
GCaMP6s (ΔF/F %)
20
5
-10
Freq (Hz)
30
20
10
0
Theta/ Delta
20
10
0
EMG
200
100
0
Delta
6000
3000
0
0
200
400
600
800
1000
Time (Sec)

Figure S8

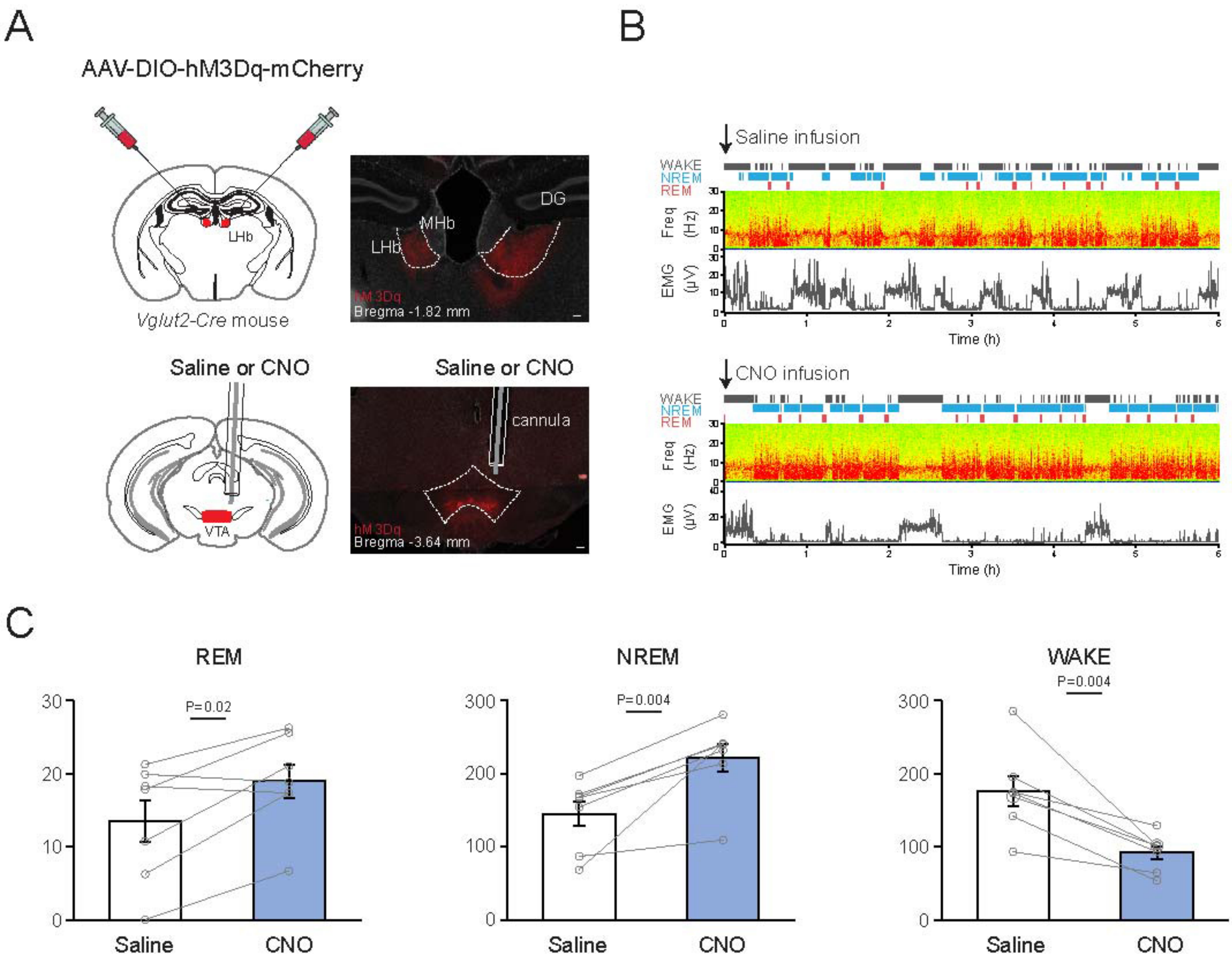

A
AAV-DIO-hM3Dq-mCherry
LHb
Vglut2-Cre mouse
MHb
DG
LHb
hM3Dq
Bregma -1.82 mm
Saline or CNO
VTA
Saline or CNO
cannula
hM3Dq
Bregma -3.64 mm
B
Saline infusion
WAKE
NREM
REM
Freq (Hz)
EMG (µV)
Time (h)
CNO infusion
WAKE
NREM
REM
Freq (Hz)
EMG (µV)
Time (h)
C
REM
P=0.02
Saline
CNO
NREM
P=0.004
Saline
CNO
WAKE
P=0.004
Saline
CNO


Figure S9

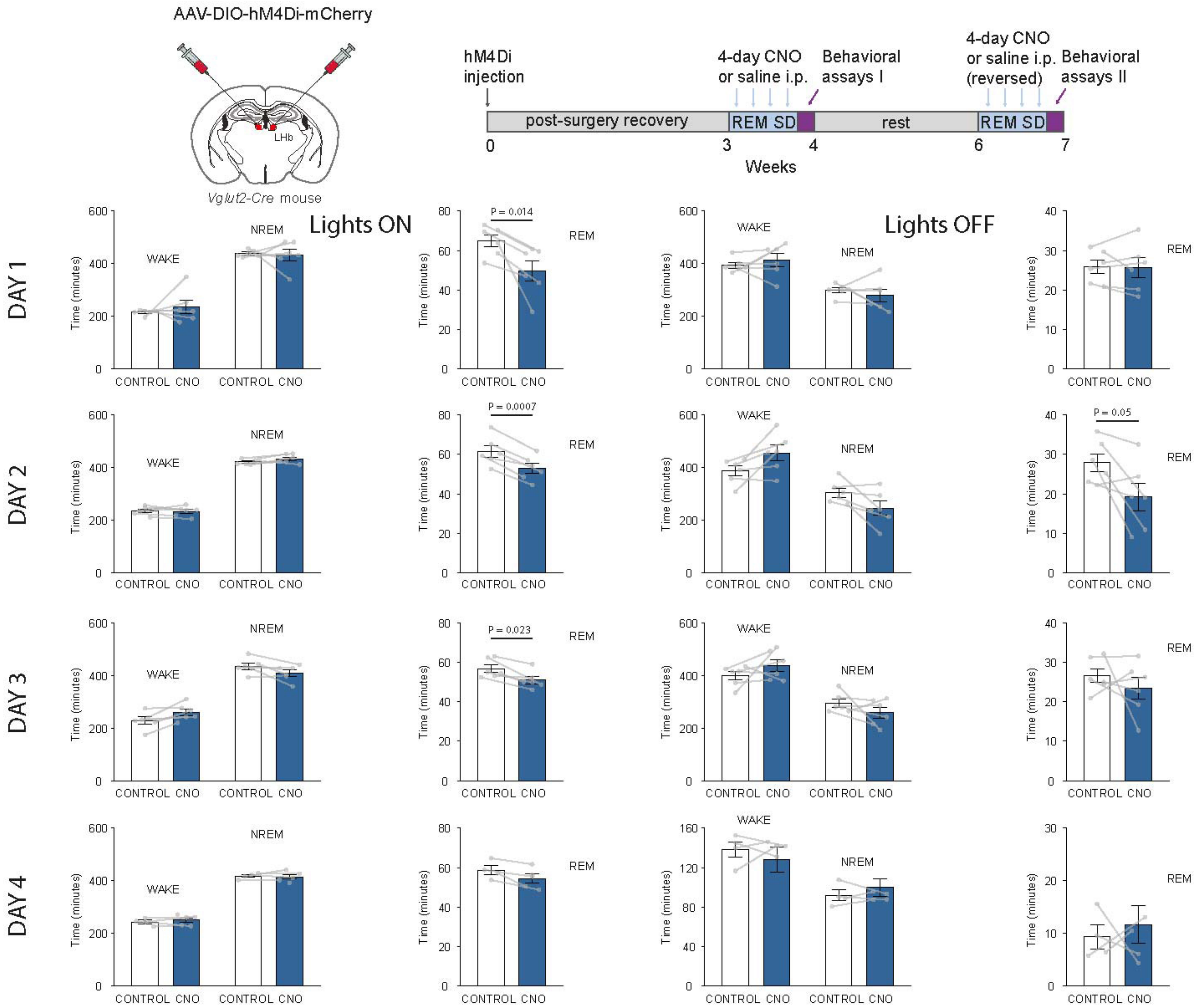
AAV-DIO-hM4Di-mCherry
LHb
Vglut2-Cre mouse
hM4Di injection
4-day CNO or saline i.p.
Behavioral assays I
4-day CNO or saline i.p. (reversed)
Behavioral assays II
post-surgery recovery
REM SD
rest
REM SD
0
3
4
6
7
Weeks
Lights ON
Lights OFF
DAY 1
DAY 2
DAY 3
DAY 4
Time (minutes)
WAKE
NREM
REM
CONTROL CNO
P = 0.014
P = 0.0007
P = 0.05
P = 0.023

Figure S10
